# GapR is a nucleoid-associated protein that stiffens and overtwists DNA to promote transcription and replication

**DOI:** 10.64898/2026.01.13.699244

**Authors:** Xinjue Wei, Lucy M. Kwiatkowski, Ryo Kawamura, John F. Marko, Monica S. Guo

## Abstract

GapR is an α-proteobacterial nucleoid-associated protein reported to either directly regulate transcription of AT-rich DNA or to regulate transcription indirectly through sensing DNA topology and modulating topoisomerase activity. We use single-DNA micromechanics, biolayer interferometry (BLI), and computational analysis to study GapR transcriptional regulation. Micromechanics experiments show that GapR overtwists DNA and shortens its contour length. GapR binding also substantially increases DNA bending persistence length and twist stiffness, while promoting duplex DNA melting at higher forces. Nonequilibrium binding experiments show GapR-DNA complexes to be extremely stable, with essentially no dissociation on hour-long time scales. Strikingly, our BLI experiments show that GapR has high affinity for AT-rich DNA while our micromechanics show that GapR binding is enhanced by pre-twisting of DNA, validating GapR affinity for both forms. By analyzing published GapR binding data, we differentiate between AT-rich and overtwisted DNA binding, revealing that overtwisted DNA primarily determines GapR localization. Lastly, we show that these two binding modes exert opposite effects on transcription. AT-rich binding stimulates initiation, likely by promoting RNA polymerase progression, while overtwisted DNA binding represses transcription, possibly by stabilizing local overtwist at promoters. We propose that GapR binding promotes a conformation of DNA ideal for efficient replication and transcription progression.

**Key points:**

- GapR is a DNA structuring protein that binds AT-rich and overtwisted, positively supercoiled DNA
- GapR binding stiffens DNA, limits plectoneme formation, and melts undertwisted DNA at higher force
- GapR binding modes exert opposing effects on transcription initiation while promoting elongation

## Introduction

Bacterial chromosomes must be compacted by three orders of magnitude to fit inside a cell. To accomplish this feat, DNA is organized and compacted by supercoiling, and nucleoid-associated proteins (NAPs) (1–4). Supercoiling is either “negative” or “positive”, with both types generating changes in DNA twist and DNA crossings (writhes). Negative supercoils (-SC) undertwist DNA, forming a left-handed superhelix, but with the unwinding torsional stress promoting DNA duplex separation. -SC promotes transcription and replication initiation and elongation (5, 6). Positive supercoils (+SC) overtwist DNA, forming a right-handed superhelix that generates plectonemes under tension. +SC suppresses strand separation, and blocks transcription and replication (5, 6). Both replication and transcription generate supercoiling. Replication produces +SC in front of the replisome while transcription follows the ‘twin-domain model’ and generates +SC ahead of RNA polymerase (RNAP), leaving -SC in its wake (6–8). Essential enzymes called topoisomerases are required to break and rejoin the DNA duplex to remove supercoiling, enabling progression of DNA and RNA polymerase (9, 10).

NAPs are a diverse group of small, basic proteins that condense DNA by bending, bridging, filamenting, or wrapping it (2, 11). Although NAPs are architectural proteins, NAPs can also be pleiotropic transcriptional regulators (5) . How NAP binding impacts transcription is poorly understood for three reasons. First, NAPs restructure the DNA double helix, altering the biophysical parameters of the DNA (e.g. stiffness, and contour length) (12–15). Second, NAPs recognize both DNA sequence and supercoiling (5, 11) , potentially altering local DNA topology. Lastly, where a NAP binds in a transcription unit impacts regulation. Therefore, NAP function is an emergent behavior dependent on how the NAP structures DNA, whether binding traps supercoiling, and how these activities in genomic context block or activate RNA polymerase. Even for well-studied NAPs such as *Escherichia coli* HU, which stimulates or represses >1000 genes (16) , how each binding event causes a particular transcriptional outcome remains unclear.

To better understand how NAPs regulate transcription, we must combine a biophysical, topological, and genome-wide understanding of NAP function. A good NAP to study using this framework is α-proteobacterial GapR, as it has been reported to be both a transcriptional regulator that binds AT-rich DNA and a +SC sensor that promotes replication and regulates transcription indirectly through modulating topoisomerase activity (17–20). Here, we combine biophysical and topological studies with an analysis of GapR binding and its impact on transcription to clarify whether GapR is a transcriptional regulator.

GapR was initially identified as an essential DNA-binding protein that recognizes AT-rich DNA to regulate gene expression in *Caulobacter crescentus* (17–19). Our subsequent structural and biochemical work demonstrated that GapR forms tetrameric complexes that encircle overtwisted, +SC DNA (20). We also showed that GapR stimulates +SC removal by topoisomerases *in vitro* and is consequently essential for replication progression *in vivo* (20). Our studies of GapR localization with chromatin immunoprecipitation sequencing (ChIP-seq) and others’ microscopy work validated this idea, showing GapR binds ahead of replication forks and at highly-expressed transcription units (19–21). While our ChIP-seq captured GapR binding to both AT-rich and +SC DNA, we did not observe correlation between binding and local transcriptional activity, leading to our proposal that GapR is not a transcriptional regulator but a topological sensor that marks +SC for topoisomerase removal (20).

More recent work supports both AT- and +SC-dependent binding modes and a role in transcription. While our initial GapR crystal structure suggested binding to +SC DNA, further structural studies showed binding to AT-rich, non-supercoiled DNA (22, 23). Additional support for potential +SC binding comes from our single-molecule magnetic tweezer (MT) studies, which demonstrated that GapR binding overtwists DNA (21). Lastly, atomic force microscopy experiments suggested that GapR oligomerizes and bridges DNA like H-NS (24), implying that binding may silence transcription. Taken together, these data leave the function of GapR incompletely understood.

Here, we show that GapR has a secondary function in regulating transcription in addition to its primary role in replication. We find that GapR binds both AT-rich and +SC DNA, with these two binding modes having opposite effects on transcription. First, using MT and biolayer interferometry (BLI) experiments, we provide detailed biophysical and topological analysis of GapR-DNA interactions. We demonstrate that GapR binds to and overtwists DNA, increasing its stiffness, and facilitates strand separation under torsional stress, particularly in AT-rich sequences. We also find that GapR has higher affinity for AT-rich DNA than for GC-rich DNA, and that binding is enhanced on pre-overtwisted DNA. Next, using published ChIP-seq and RNA-seq datasets, we separate AT-and +SC-dependent GapR binding, revealing that +SC-dependent binding is the major determinant for GapR localization during growth. We observe that AT-rich and +SC-dependent binding occur at separate sites within a transcription unit and have opposite effects on local promoter activity. We further find that both binding modes support RNAP progression. Together, we propose that GapR holds DNA in a rigid overtwisted confirmation, supporting efficient DNA loading into replication and transcription complexes.

## Results

### GapR positively shifts DNA extension versus linking number response

To better understand the biophysical and topological properties of GapR, we examined GapR using MT, an assay that interrogates the mechanics of GapR binding to single DNA molecules with controlled superhelical density. Briefly, one end of an ∼11.4 kb DNA fragment was immobilized to a coverslip while the other end was bound to a magnetic bead in a flow cell. The flow cell was placed in a vertical magnetic tweezer platform that allows tension (stretching force) and supercoiling to be adjusted based on the position and rotation of the magnet. At low forces (0.25 pN), rotation of the magnet introduces either over- or undertwisting into the DNA, which converts into either +SC or -SC (writhe) (**Fig. 1**).

**Fig. 1.**
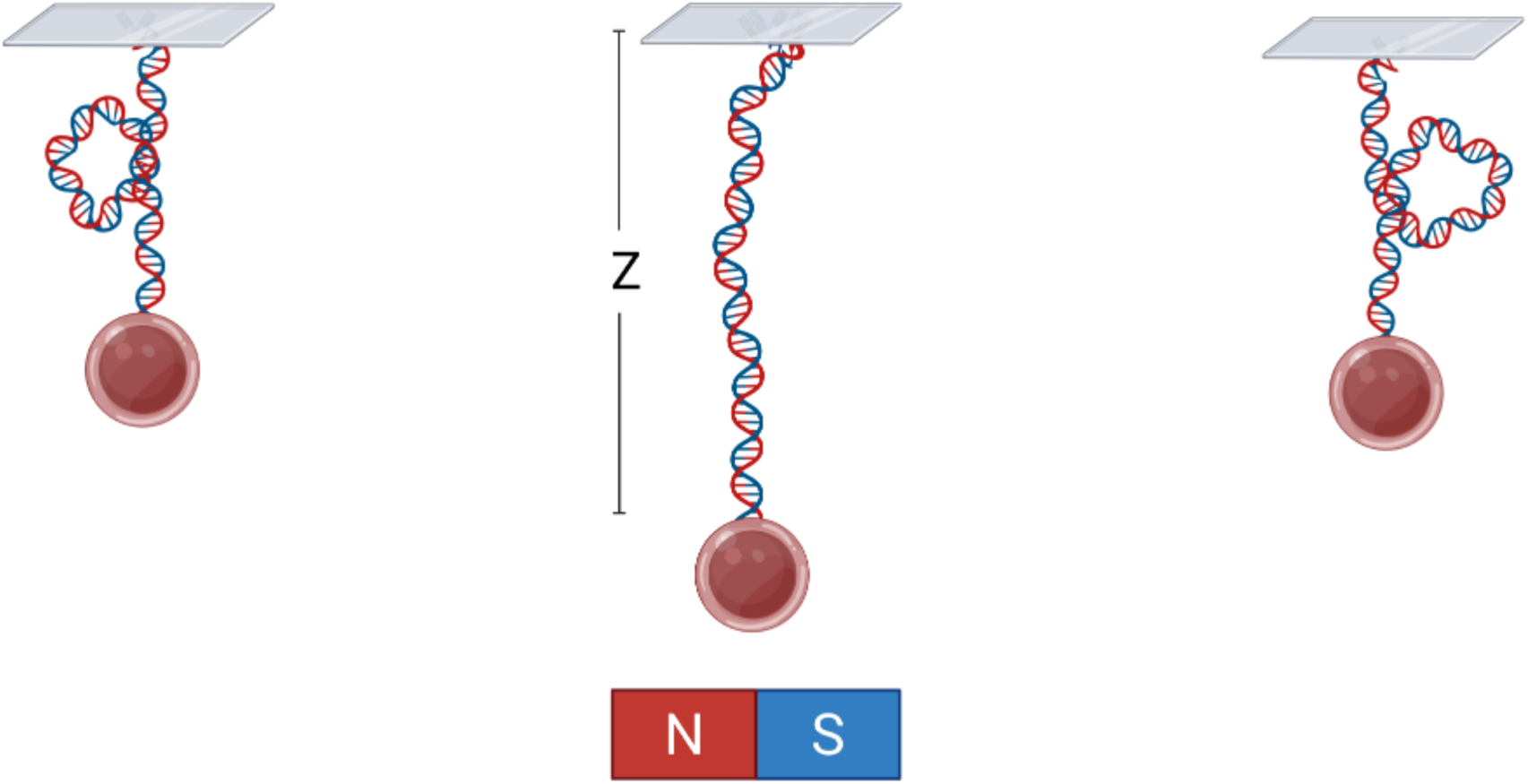
Schematic diagram of GapR-dsDNA magnetic tweezers experiments at controlled linking number and force. The double-stranded DNA (dsDNA) molecule is tethered between a 2.7 µm diameter paramagnetic bead (red), coated with streptavidin, and an anti-digoxigenin-coated glass coverslip via a biotin-labeled handle. Permanent magnets beneath the coverslip modulate the applied tension via their position. The linking number of the DNA can be altered by rotating the magnets, which rotate the bead. The middle schematic illustrates the tether in its relaxed or stretched-twisted conformation. Sufficiently large deviations from the relaxed linking number induce the formation of plectonemes, shown in the left and right panels. When GapR proteins bind to the dsDNA they influence its mechanical response (extension as a function of force and linking number).

We first assessed the relationship between GapR and DNA extension length and linking number. Fig. 2a displays a representative single-molecule DNA extension as a function of linking number under a constant tension of 0.25 pN. Traces for naked tether DNA (green), and the same DNA tether in the presence of 800 nM GapR (purple) are shown, with data acquired by a Labview program that controlled the experiment. For naked DNA, the extension increases nearly symmetrically upon underwinding (negative turns) or overwinding (positive turns), reaching a maximum of approximately 2.5 µm near zero linking number.

**Fig. 2.**
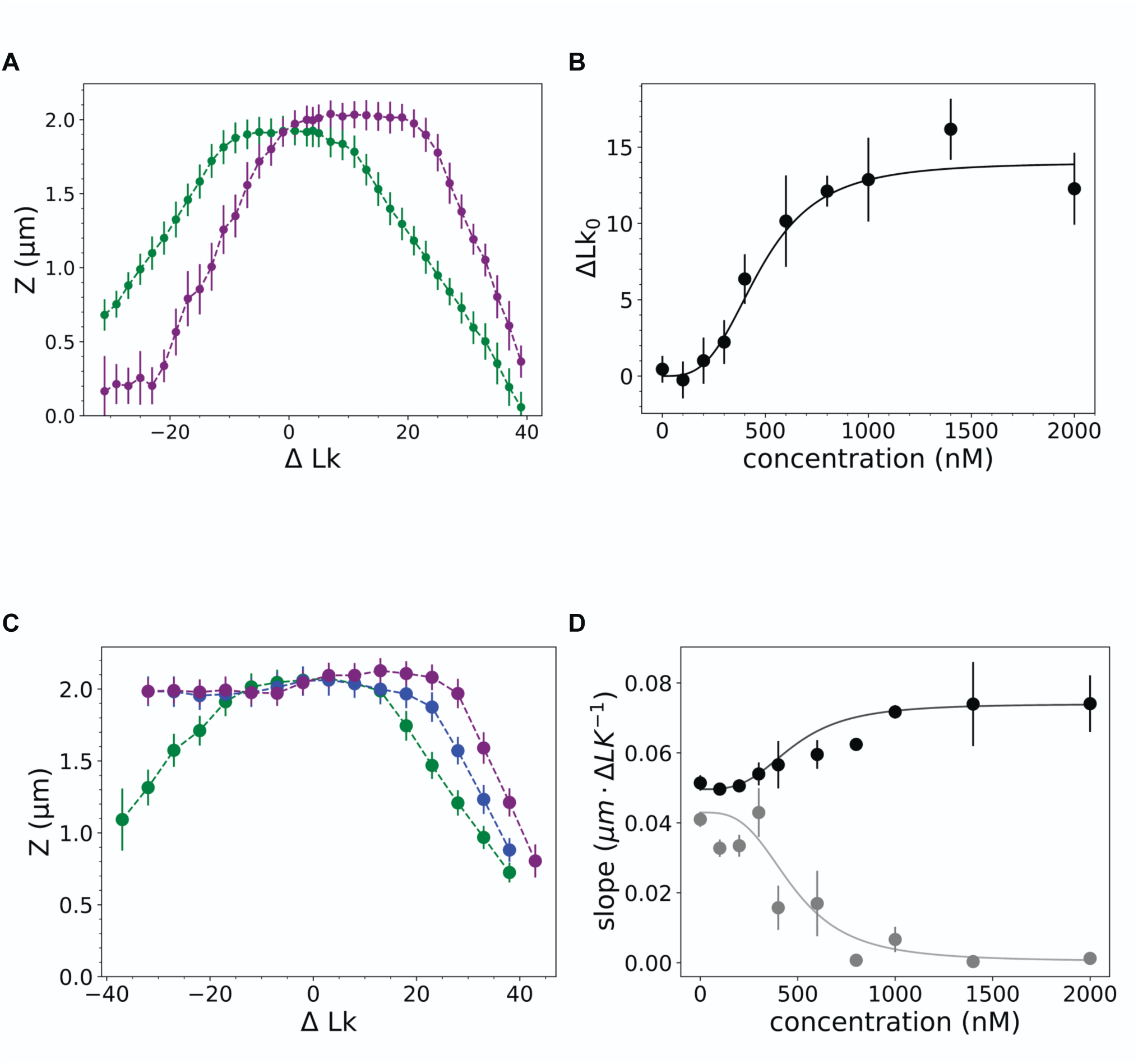
Effect of GapR on DNA extension versus linking number at fixed force. (a) GapR binding overtwists DNA. Left (green) curve: extension-linking number relation for naked DNA at 0.25 pN. Right (purple) extension-linking number relation of the same dsDNA for 800 nM GapR protein. Error bars show standard errors from time series for a single DNA, N = 250 independent extension measurements per data point. (b) Shift of relaxed linking number (*ΔLK*_0_) due to the binding of GapR protein at fixed force of 0.25 pN. Data points represent the mean ΔLk_0_ (from ≥3 independent measurements), and the solid line indicates the best-fit titration curve, corresponding to a K_d_ = 475 and a Hill coefficient of 3.25 (see text). Means and standard errors are obtained from averaging ΔLk_0_ measured in separate experiments, ranging from N =3 at high concentration, to N=9 at low concentrations. (c) GapR protein facilitates the melting of dsDNA at higher forces. Green: linking number-extension relation of naked DNA at 0.5 pN. Blue: linking number-extension relation of the same tether under 0.5 pN in 600 nM GapR solution. Purple: linking number-extension relation of the same tether under 800 nM solution. Mean and standard errors as in (a), N=300 independent measurements per data point. (d) Melting of dsDNA is correlated with the concentration of GapR in the solution. Black: the average slope of linking number-extension relation of the right plectonemic region. Gray: the average slope of linking number-extension relation of the left plectonemic region. Both solid lines indicate titration curves derived from (b) with the same Hill coefficient and K_d_. Means and standard errors were determined from separate experiments, ranging from N = 3 at high concentrations to N=9 at low concentrations.

This response reflects the characteristic behavior of torsionally constrained DNA at low force, where initial rotation decreases the extension until buckling occurs and leads to the formation of plectonemes, further reducing the extension (25, 26). The maximum extension also increases, indicating a change in both the relaxed superhelical density and the overall DNA extension under torsional stress.

Fig. 2a also shows that in the presence of 800 nM GapR, the peak of the DNA extension versus linking number curve shifts to positive linking number relative to naked DNA, indicating that GapR binding alters the DNA’s relaxed superhelical density (we use ΔLk_0_ to indicate the linking number at which the DNA extension is maximal at a given condition, with ΔLk_0_=0 for “naked” DNA; ΔLk_0_ has essentially no dependence on force over the range of forces used in this paper). There is a modest increase in extension at this concentration, indicating a change in both the relaxed superhelical density and the overall DNA extension under torsional stress.

Consolidating data from multiple extension vs. linking number measurements at various GapR concentrations, Fig. 2b illustrates the shift in ΔLk_0_ as a function of protein concentration. Data points represent the mean ΔLk from ≥ 3 independent measurements. The solid line indicates the best-fit Hill-type titration model (see Methods), yielding an apparent dissociation constant (K*_d_*) of 475 ± 30 nM and a Hill coefficient of 3.25 ± 0.35. This relatively high Hill coefficient indicates pronounced cooperativity, suggesting that the binding of one GapR molecule markedly enhances the propensity for additional GapR molecules to bind. As the protein concentration approaches 1000 nM, ΔLk_0_ plateaus, indicating that the DNA is saturated with GapR. Beyond this point, further increases in GapR concentration do not alter ΔLk_0_, signifying that the 10 kb pFos1 DNA has reached its maximal GapR binding capacity.

At a higher pulling force (remaining within a physiologically relevant range), GapR protein facilitates the melting of double-stranded DNA (dsDNA, Figure 2c). Under a tension of 0.5 pN, the linking number–extension profile for naked dsDNA tether (green), serving as a baseline, shows relatively symmetric behavior under positive and negative overtwisting. In the presence, of 600 nM (blue) and 800 nM (purple) GapR, the left portion of the extension-linking number curve, corresponding to -SC, remains flat. This indicates an unwinding of dsDNA (26–28), suggesting that GapR binding alters the torsional response of dsDNA, promoting partial melting of the double helix.

Combining results from experiments across multiple GapR concentrations shows a correlation between dsDNA melting and the concentration of GapR in solution (Fig. 2d). The black data points represent the average slope of the linking number–extension relationship for the right (overtwisted, +SC) plectonemic region, whereas the gray data points correspond to the left (undertwisted, -SC) plectonemic region. Both sets of data are shown with titration curves (solid lines) that use the Hill coefficient and dissociation constant (*K_d_*) fit in Fig. 2b.

The consistency underscores the concentration-dependent effect of GapR on dsDNA melting across underwound states and resistance to plectoneme formation in overwound states. Our data show that GapR binding overtwists DNA, acting as a local twist reservoir, and promotes unbound DNA to melt instead of forming left-handed plectonemes at forces close to those generated by physiological supercoiling (0.5 pN for σ » -0.05) (26).

### GapR binding increases bending persistence length and slightly reduces molecular length of dsDNA

Our data suggest GapR recognizes overtwisted DNA and binding promotes DNA melting under force. We next wondered how this GapR binding impacts the mechanical properties of DNA, such as length and stiffness. The random thermal motion of naked DNA follows the worm-like chain model. Upon protein binding DNA segments frequently stiffen and behave as rodlike elements over defined lengths. This rodlike behavior is quantified as the bending persistence length, and can be determined by relating the applied force on GapR-bound DNA to the maximum extension across a series of GapR concentrations. Fig. 3a shows a representative force–extension curve for a nicked DNA tether (ΔLk = 0) as a function of pulling force. Traces for naked DNA (green), as well as the same tether in the presence of 600 nM (purple) and 800 nM (blue) GapR are shown. Fig. 3b provides a transformed view of these data, with 1/√F on the x-axis. In this representation, the y-intercept corresponds to the DNA contour length, while the slope reflects the persistence length consistent with the worm-like chain model (29). As GapR concentration increases, the slope becomes progressively larger — smallest for naked DNA, intermediate at 600 nM and steepest at 1400 nM. This indicates an increase in DNA stiffness with increased GapR. In contrast, the contour length remains relatively constant, exhibiting only a minor reduction as GapR concentration rises.

**Fig. 3.**
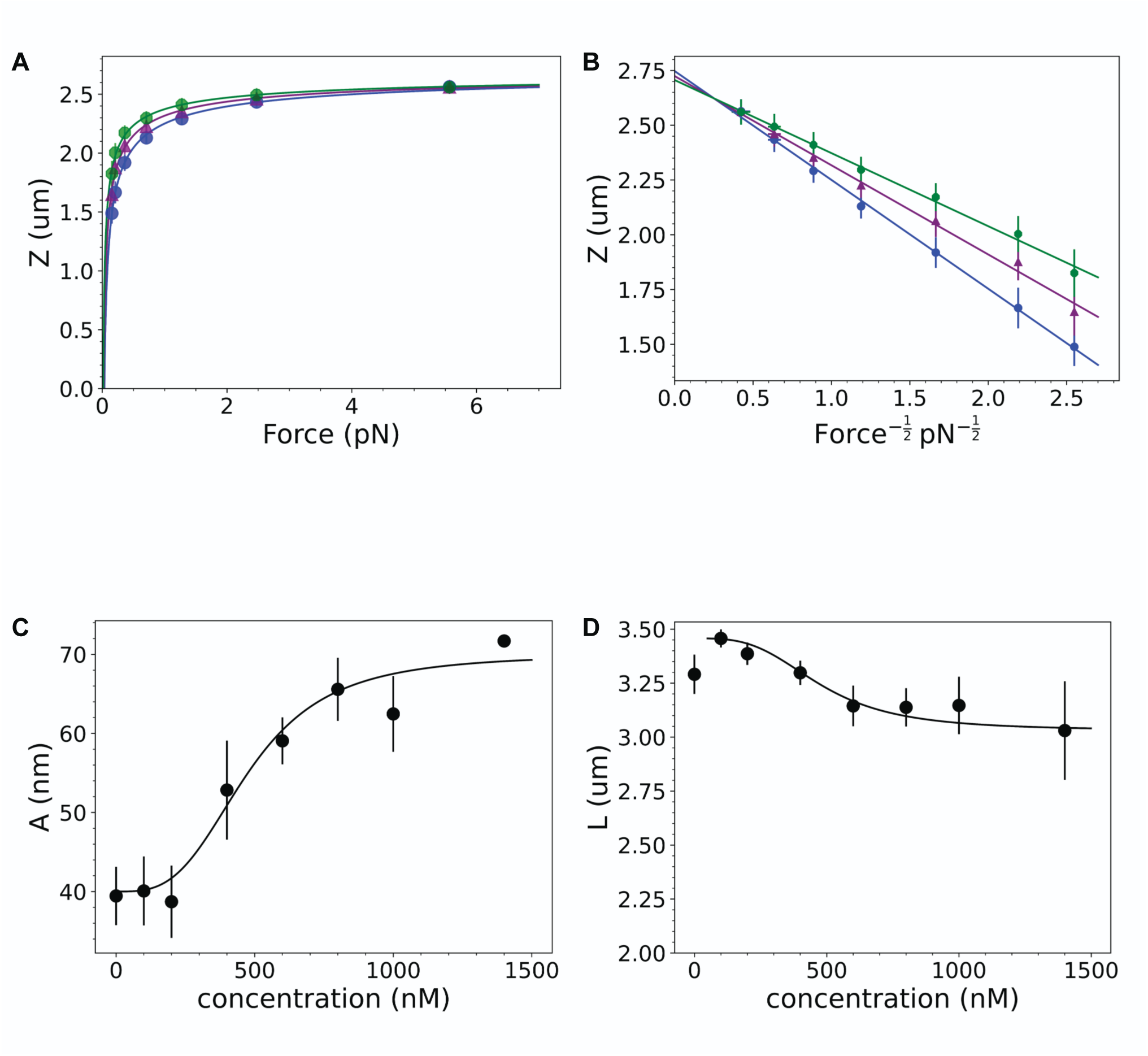
GapR protein binding alters bending persistence length and molecular length of dsDNA. (a) GapR binding changes the force-extension relation of dsDNA. Green curve: force-extension relation of naked DNA. Purple: force-extension of the same tether in 600 nM GapR. Blue: Purple: force-extension of the same tether in 1400 nM GapR. Mean and standard errors as in Fig 2(a), N=50 independent measurements per data point. (b) Linearized plot of the force-extension curve. The y-intercept indicates contour length; slope is related to persistence length (see text). Standard errors are the same as those of (a). (c) GapR increases the persistence length of dsDNA. Data points represent the mean persistence length (from ≥3 independent measurements), and the solid line indicates the best-fit titration curve from Fig. 2(b). Means and standard errors were determined from separate experiments, ranging from N = 3 at high concentrations to N =8 at low concentrations. (d) GapR decreases the contour length of dsDNA. Data points represent the mean contour length (from ≥3 independent measurements). The solid line indicates the best-fit titration curve from Fig. 2 (b). Means and standard errors were determined from separate experiments, ranging from N = 3 at high concentrations to N =7 at low concentrations.

Experiments performed at multiple GapR concentrations, with four or more independent experiments in each case, yielded the data presented in Fig. 3c. These measurements illustrate a pronounced increase in bending persistence length as GapR binding sites become progressively occupied. At lower concentrations (≤250 nM), the persistence length remains near 40 nm, in agreement with values expected for naked dsDNA (29–32).

Beyond this threshold, the DNA stiffens considerably, ultimately reaching ∼70 nm at saturating levels (1250–1500 nM). This marked stiffening behavior is captured by the titration fit of Fig. 2b, shown in black, emphasizing the strong concentration-dependent influence of GapR on DNA rigidity.

Analyzing the contour length from the worm-like chain fit shows that the contour length of the DNA-GapR tether decreases with increasing GapR concentration (Fig. 3d), albeit to a lesser degree than the observed changes in persistence length. This indicates a subtle DNA compaction phenotype in response to GapR binding. The decreased contour length again closely follows the titration relationship of the binding curve. This observation explains the net increase in tether extension: although the persistence length nearly doubles, the overall tether extension rises by only ∼15%, as the reduction in contour length offsets some of the gain resulting from the increased bending persistence length (mechanical stiffness). Together, these data highlight the impact GapR binding has on the DNA stiffness, showing an increased bending persistence length. GapR binding does not greatly increase overall DNA extension since the additional length expected from the increased persistence length is partially offset by the localized DNA compaction.

### GapR binding increases DNA torsional rigidity

Torsional stiffness is a fundamental mechanical parameter of dsDNA, as described by the twistable worm-like chain (TWLC) model, and is related to the curvature of the extension versus linking number relationship near its peak (near ΔLk_0_), prior to the onset of plectoneme formation (26, 28, 33, 34). Fig. 4a shows the curvature of the extension versus linking number curve at different GapR concentrations. Under applied forces of 0.25 pN (green) and 0.5 pN (blue), this curvature remains nearly constant across the range of 0 – 1400 nM. Curvature data for 0.5 pN with GapR concentration higher than 300 nM is absent due to extensive melting (Fig. 2d); the resulting flat response makes accurate determination of the curvature problematic (and inappropriate to analyze using the simple TWLC due to the DNA being a mixture of strand-separated and base-paired regions).

**Fig. 4.**
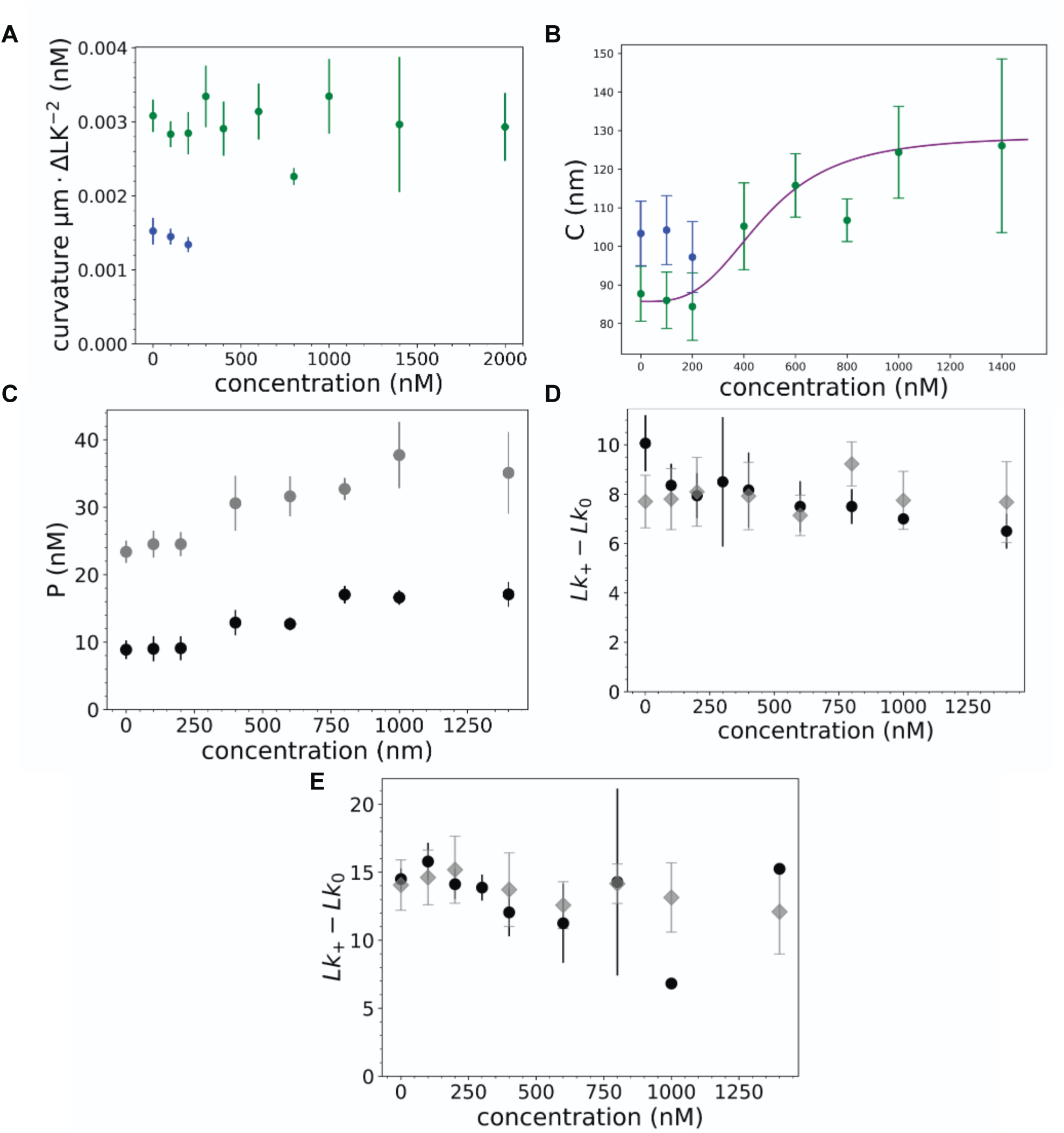
Effect of GapR binding on DNA torsional rigidity. (a) Curvature of the linking number-extension relation (pre-buckling) is unaffected by GapR protein. Top (green): The curvature of linking number-extension relation at different GapR concentrations at 0.25 pN. Bottom (blue): The curvature at 0.5 pN. Means and standard errors were determined from separate experiments, ranging from N = 5 at high concentrations to N =8 at low concentrations. (b) GapR increases the twist persistence length of dsDNA. Green: twist persistence length of dsDNA calculated from curvature at 0.25 pN. Purple: best-fit titration curve from Fig. 2(b). Blue: twist persistence length of dsDNA calculated from curvature at 0.5 pN. Means and standard errors were determined from separate experiments, ranging from N = 3 at high concentrations to N =7 at low concentrations. (c) GapR protein binding slightly increases the plectonemic twist persistence length, indicating an increase in the energy required for dsDNA to absorb supercoiling into plectonemic structures. Gray: plectonemic twist persistence length calculated with extension-linking number curve at 0.5 pN. Black: plectonemic twist persistence length calculated with extension-linking number curve at 0.25 pN. Means and standard errors were determined from separate experiments, ranging from N = 3 at high concentrations to N =7 at low concentrations. (d) GapR protein binding slightly decreases the relative energy barrier for plectoneme formation at 0.25 pN, as indicated by reductions in the buckling width—the difference in linking number (ΔLk) between the positive buckling point and Lk_0_ (see main text for details). Black: observed buckling width. Gray: expected buckling width calculated from the twistable worm-like-chain model based on DNA mechanical properties (results of Figs. 2 and 3). Means and standard errors were determined from separate experiments, ranging from N = 3 at high concentrations to N =7 at low concentrations. (e) GapR protein have slightly decrease the relative energy barrier of forming a plectoneme at 0.5 pN, again reflected by the reduced buckling widths. Black: observed buckling width. Gray: expected buckling width calculated from the twistable worm-like-chain model based on DNA mechanical properties (results of Figs. 2 and 3). Means and standard errors were determined from separate experiments, ranging from N = 3 at high concentrations to N =7 at low concentrations.

Integrating the curvature measurements with the persistence and contour lengths obtained in Figs. 3c and 3d, the twist persistence length (or twist stiffness in *k_B_T* energy units) of dsDNA was determined (see Methods). As shown in Fig. 4b, increasing GapR concentration leads to a marked enhancement in torsional rigidity, with data acquired at 0.25 pN (green) and 0.5 pN (blue, for GapR concentration lower than 300 nM). Upon saturation of binding sites at concentrations above 1000 nM, the twist persistence length increases by over 30%, rising from 90 nm to 120 nm. Notably, this enhancement closely follows the titration fit derived from the GapR binding curve (Fig. 2b, purple), underscoring the robust, concentration-dependent modulation of DNA torsional stiffness by GapR.

Another important aspect of DNA torsional rigidity arises from the torsional stiffness of the plectoneme. The plectonemic torsional persistence length (*P*, conceptually similar to a twist persistence length for a straight rod) (26, 35) quantifies the energy cost associated with introducing supercoils into the plectoneme. This parameter is expected to be force dependent as it is based on a quadratic approximation for a more complex underlying plectoneme free energy (26). As illustrated in Fig. 4c, GapR binding results in a slight increase in *P* under both 0.25 pN and 0.5 pN conditions. The results for twist and plectoneme torsional stiffnesses indicate that the GapR “coating” on DNA stabilizes the double helix against torsional stress by simultaneously increasing resistance to both initial twisting and subsequent formation of plectonemic supercoils.

Despite these concurrent changes in twist and plectonemic torsional persistence lengths (stiffnesses), the buckling point remains relatively constant. The parameters *P* and *C* exert opposing effects on the ability of dsDNA to accommodate linking number changes.

Specifically, an increased *C* implies a higher energetic cost to induce torsional deformation into the DNA itself (the added stiffness from the protein sheath), while the increased *P* indicates a greater energy cost of soaking up linking number in plectonemic turns (likely due to the increased thickness from the GapR sheath).

Fig. 4d presents both measured buckling points derived directly from extension versus linking number curves (black points) and predicted buckling points (gray points) calculated using the previously determined persistence length, curvature, applied force, and the post-buckling slope of the linking number–extension relationship (see Methods). The predicted and measured buckling points at 0.25 pN are in accord, each being about 8 turns (Fig. 4d), indicating the self-consistency of the theory used for our fitting (26). A similar correspondence is evident at 0.5 pN, as shown in Fig. 4e, where both experimental and theoretical values indicate a stable buckling threshold of approximately 15 turns. The elevated buckling point at higher forces reflects an increased energetic barrier to incorporating twist into the plectoneme, leading to reduced DNA shortening prior to buckling.

### GapR shows sequence-dependent binding, with a preference for AT-rich DNA

While our micromechanics experiments support a model for GapR recognizing overtwisted DNA, we sought to directly measure GapR affinity for AT-rich and overtwisted DNA. First, we measured affinity for DNA with different AT-content using biolayer interferometry (BLI), a standard method of determining protein binding and dissociation rates. Fig. 5a directly compares the BLI-derived binding curve (green), measured with a 50 base pair DNA substrate containing a 1:1 A–T to G–C sequence ratio, to the MT data (purple) obtained using pFOS1 DNA. Notably, the BLI titration exhibits a lower Hill coefficient of 2.2 ± 0.13 and a reduced *K_d_* of 344 ± 34 nM relative to the values derived from the magnetic tweezer experiments (3.25 and 475 nM). This indicates that with increasing concentration, initial binding of GapR slightly precedes its mechanical distortion of DNA, the latter read out by the MT experiments; also this may be a result of the MT experiments probing binding to an extended DNA rather than to a short fragment under zero tension.

**Fig. 5.**
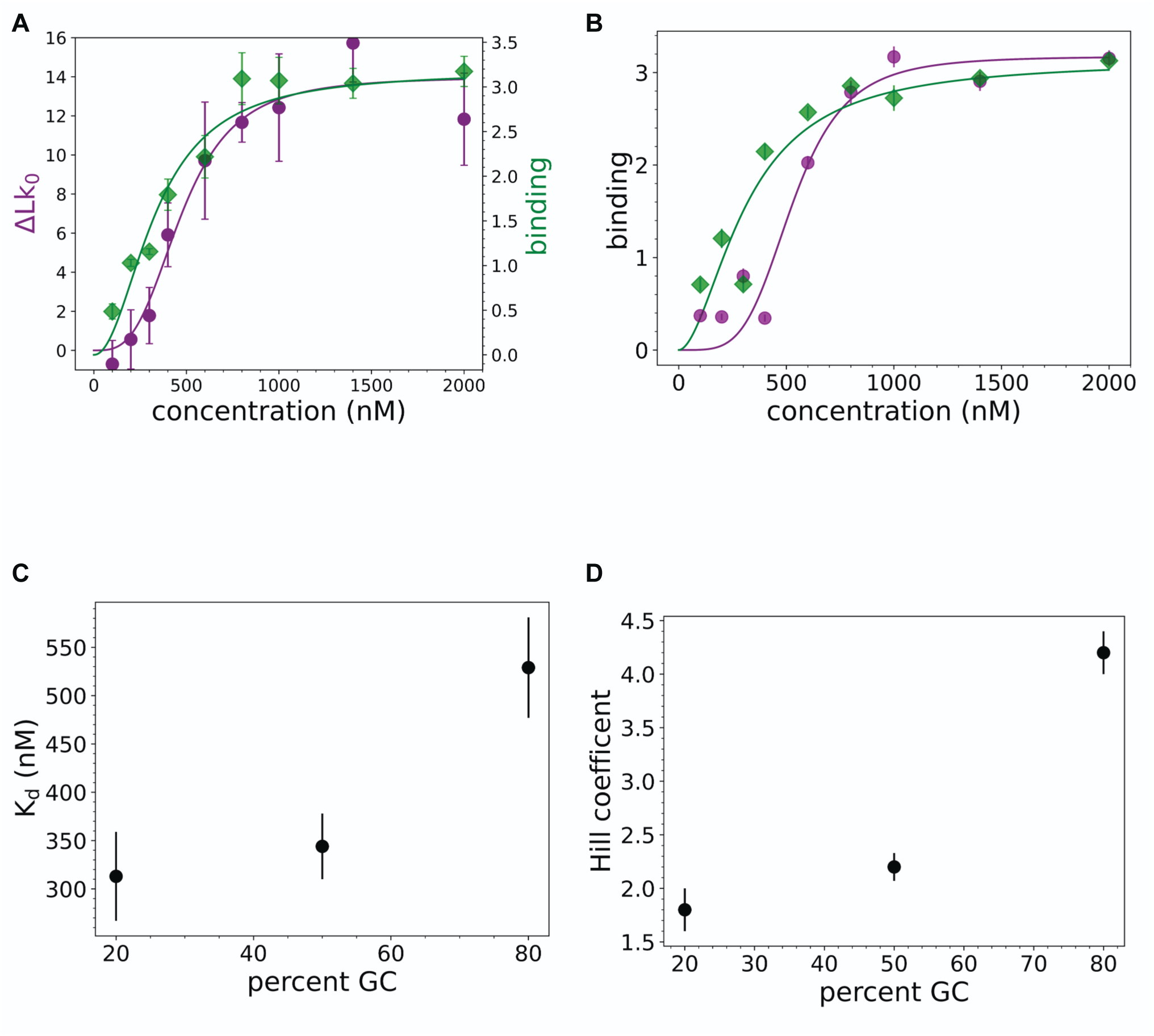
GapR binding to DNAs of varied sequence composition. (a) Binding rate measurement with biolayer interferometry (BLI). Green: binding results from BLI. Data points represent the mean signal (N=3 independent measurements), and the solid line indicates the best-fit titration curve. Purple: binding measurements from magnetic tweezer experiments. (b) GapR binding is sequence dependent. Left (green): Binding signal for A-T rich sequence in response to changes in GapR concentration. Purple: binding signal of G-C rich sequence as a response to GapR concentration. N=3. (c) GapR binding prefers a sequence with a high A-T composition. A-T rich sequence (20% G-C) have lower K_d_ compared to G-C rich sequence (80% G-C) and balanced composition sequence (50% G-C). N=3. (d) Hill coefficient follows same pattern as K_d_, with A-T rich sequence having the lowest Hill coefficient, balanced larger, and G-C rich the largest. N=3.

To investigate the DNA-sequence dependence of GapR binding, additional BLI experiments were conducted using 50 bp dsDNA segments with either 20 % or 80 % GC content (Fig. 5b; see Methods for DNA sequences used). The results reveal that the GC-rich sequence exhibits approximately twofold greater cooperativity, with a Hill coefficient of 4.2 ± 0.2, compared to 1.8 ± 0.2 for the AT-rich sequence. Furthermore, the GC-rich substrate displays a higher dissociation constant (*K_d_* = 529 ± 52) relative to that of the AT-rich sequence (*K_d_* = 313 ± 46). Figs. 5c and 5d illustrate the correlations between Hill coefficients and *K_d_* values as a function of sequence composition. These findings suggest that GapR is more likely to bind available AT-rich regions before GC-rich regions and may not require a fully formed tetramer for AT-rich binding. Then as the AT-rich regions become saturated, GapR binding extends to GC-rich regions in a fashion characterized by enhanced cooperativity.

### GapR-DNA complexes are extremely stable in protein-free solution

Next, we sought to measure GapR affinity for overtwisted DNA with MT. To do so, we first measured GapR dissociation rates, as a slow dissociation is required to observe whether pre-twisting DNA promotes additional GapR assembly. GapR dissociation does not conform to a classical binding–unbinding equilibrium where on- and off-rates are comparable near the apparent *K_d_* (36), similar to effects seen for other protein-DNA interactions (12, 37–40). To quantitatively assess this behavior, wash-out (dissociation) experiments were performed by washing the flow cell with 150 μL of GapR-free buffer (Tris with 100 mM NaCl) and incubating the GapR-bound DNA tether for 1 hour. Extension-linking number measurements were taken both before and after washing to evaluate whether sufficient GapR dissociation has occurred to restore the tether’s mechanical properties to those of the naked DNA.

Fig. 6a presents representative force–extension curves from a wash-out experiment performed with 600 nM bacterial GapR under a constant tension of 0.25 pN. The green trace corresponds to naked DNA, the purple trace to the DNA tether in the presence of 600 nM GapR, and the blue trace to the same tether after washing with 5x the flow cell volume of GapR-free solution plus one hour incubation in GapR-free solution. The peak positions and tether extensions remain nearly unchanged after the hour-long incubation in protein-free solution, indicating little or no GapR dissociation.

**Fig. 6.**
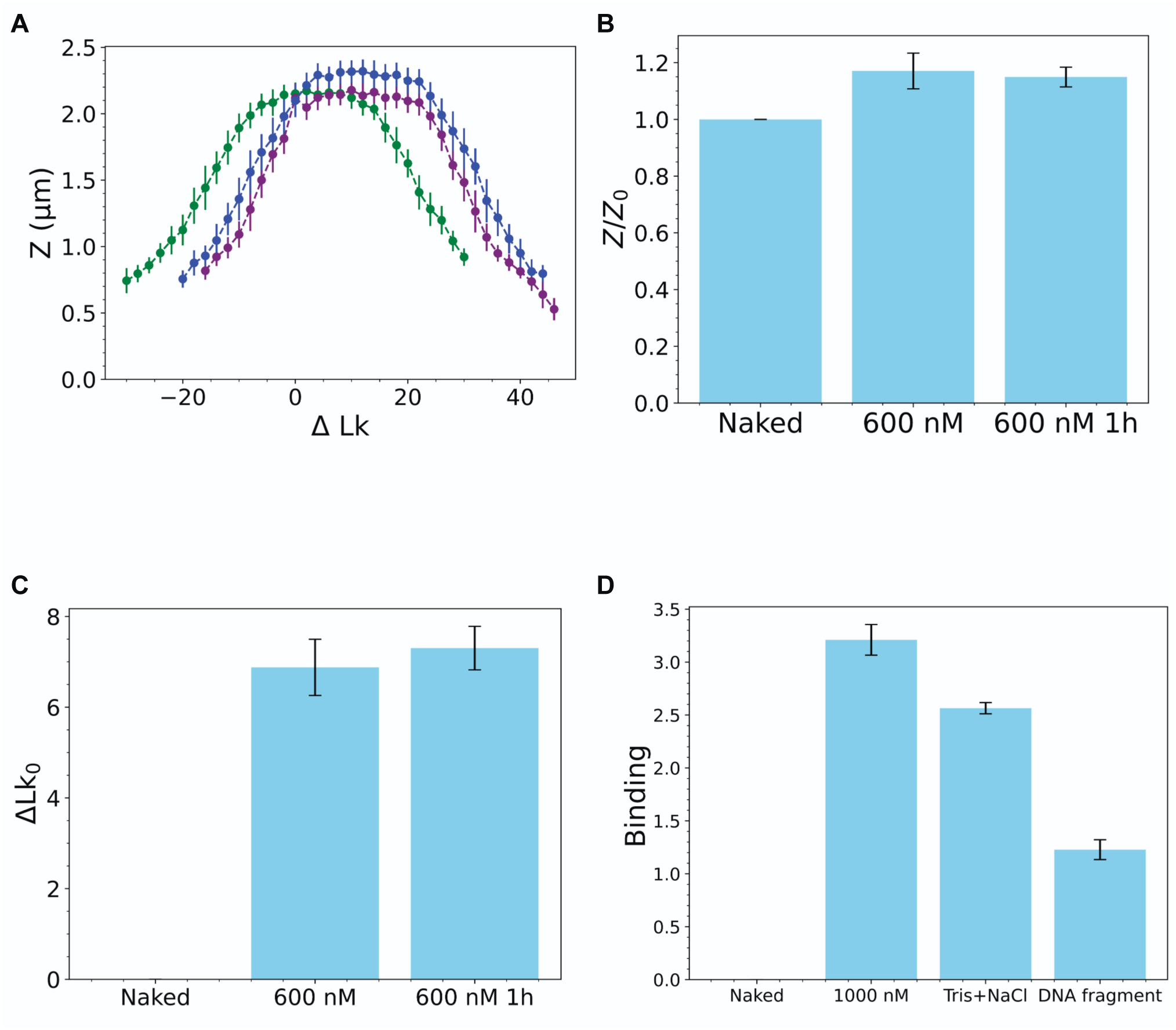
Unbinding kinetics of GapR from dsDNA to protein-free solution. (a) GapR dissociation is slow under physiological conditions. Left (green) curve: extension-linking number relation for naked DNA. Purple: the extension-linking number relation before washing. Blue: extension-linking number curve of the dsDNA tether 1 hour after removal of GapR solution. Standard error bars are computed as for Fig. 2a, with N = 250. (b) Comparison of the extension ratio at ΔLk = 20 for the naked dsDNA and the same tether in 600 nM of GapR before (middle) and after (right) washing with buffer (zero GapR concentration) for 1 hour. N=3 separate experiments were used to determine standard errors; P = 0.04 for naked vs. 600 nM; P = 0.02 for naked vs. 1 hr wash; P = 0.8 (no effect) for 600 nM vs. 1 hr wash. (c) Comparison of relaxed linking number relative to naked DNA (left) to that of the same tether in 600 nM of GapR before (middle) and after (right) washing. N = 3; P = 0.003 for naked vs. 600 nM; P = 0.0014 for 600 nM vs. 1 hr wash; P = 0.6 (no effect) for 600 nM vs. 1 hr wash. (d) Comparison of GapR protein dissociation behavior in different buffers over a 20- minute timescale, as assessed by BLI. The BLI signal obtained at a GapR concentration of 1000 nM (middle left) serves as a reference. Middle right: minimal GapR dissociation is observed in reaction buffer containing Tris and 100 mM NaCl. Right: significantly increased dissociation is observed in the presence of competing herring sperm DNA. N = 3; P = 0.05 for initial 1000 nM binding (left bar) vs. buffer-only wash (middle bar); P = 0.006 for initial 1000 nM vs. wash by DNA fragments (right bar); P = 0.0032 for buffer wash vs. DNA fragment wash.

Averaged results from three independent trials (Fig. 6b) confirm that the extension ratio (Z/Z_0_) remains constant before and after washing, and +SC of dsDNA (shift of extension versus linking number response) is retained upon removal of GapR from buffer (Fig. 6c). BLI experiments (Fig. 6d) also showed a minor change in binding signal of GapR in Tris-100 mM NaCl solution; a relatively large binding signal reduction occurred when 10 ng/uL herring sperm DNA fragments were introduced, indicative of competitive unbinding (transfer of GapR to the herring DNA).

### GapR preferentially binds to overtwisted dsDNA

Given that GapR induces overtwisting in dsDNA (Fig. 2) and shows very slow unbinding kinetics (Fig. 6), we carried out experiments to test how DNA conformation impacts GapR binding. DNA tethers were first incubated with GapR under torsionally relaxed conditions (Lk = 0) to facilitate binding. Unbound protein was removed by exchanging the chamber with 150 *μL* protein-free buffer, and extension–linking number curves were to probe the mechanical response. +SC were then introduced to impose torsional stress, GapR was re-introduced under these overwound conditions, and excess protein was again washed out. Repeating the measurements under the same force regimen enabled direct comparison of GapR–DNA interactions in relaxed versus overwound states.

Fig. 7a compares the linking number–extension curves for naked DNA (green), with the result obtained for GapR binding at ΔLk = 0 (purple) followed by washout of protein solution (washout occurred before linking number was changed). As expected, the binding of GapR leads to a shift of the peak of the curves towards a larger linking number. Then linking number was changed to ΔLk = 15 (blue) and 400 nM GapR solution was added. Again, the protein solution was washed out, and the extension as a function of linking number was measured. The resulting curve for binding at ΔLk = 15 is shifted toward higher linking numbers compared to that for ΔLk = 0 condition. Averaged results across four experiments demonstrate a clear preference of GapR for +SC dsDNA, as illustrated in Fig. 7b.

**Fig. 7.**
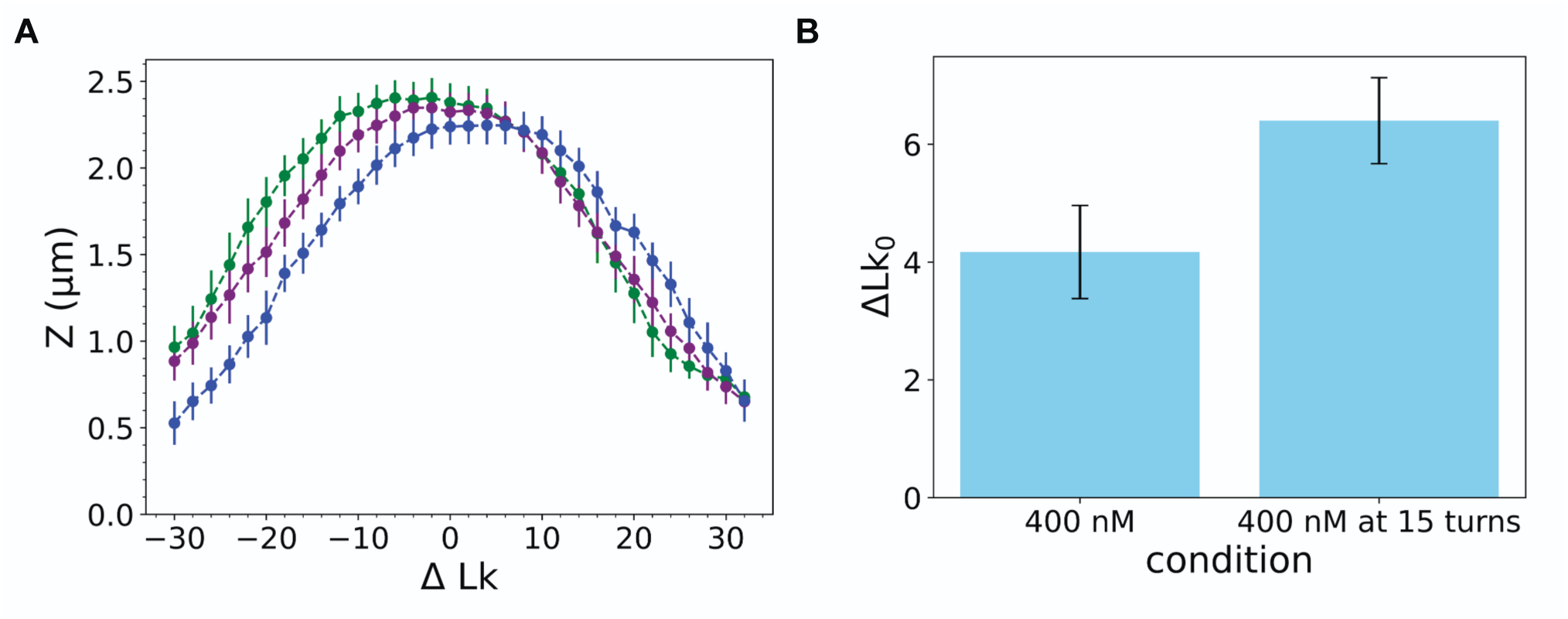
GapR exhibits increased binding affinity for positively overtwisted dsDNA. (a) An experiment comparing extension versus linking number curve of DNA tether in the absence of GapR (green), following GapR binding at 400 nM with ΔLk = 0 (purple), and following GapR binding at 400 nM with ΔLk = 15 (blue). Standard errors determined as in Fig. 2(a) for N = 130. (b) Average shifts in Lk_0_ from four independent experiments. Left: average Lk_0_ shift upon binding at ΔLk = 0; right: average Lk_0_ shift upon binding at ΔLk = 15. P = 0.13.

### Transcriptional inhibition isolates AT-dependent GapR binding

Previous *in vivo* studies on GapR binding have noted its association with either AT-rich or both AT-rich and +SC DNA (17–20). As our biophysical data show both binding modes, we sought experimental conditions to differentiate them. We reasoned that treatment with rifampicin (rif), an RNA polymerase inhibitor (41), would eliminate transcription-mediated supercoiling (in particular the +SC ahead and -SC behind RNAP), allowing us to examine GapR behavior in the absence of transcription-generated SC. We therefore analyzed published GapR ChIP-seq in untreated (GapR) and rif-treated (GapR+rif) cells (20).

First, we visually compared GapR profiles to the local AT content. While the untreated GapR profile frequently deviated from the local AT content, the GapR+rif profiles better reflected it (Fig. 8a, S2a). To quantitate how well GapR binding correlates with AT content genome-wide, we calculated the covariance between the local AT content and each profile in 10 kb sliding windows (Fig. 8b). While GapR binding is well-correlated with AT-richness (mean Pearson’s *r* = 0.581), correlation notably increases upon rif treatment (mean Pearson’s *r* = 0.721).

**Fig. 8.**
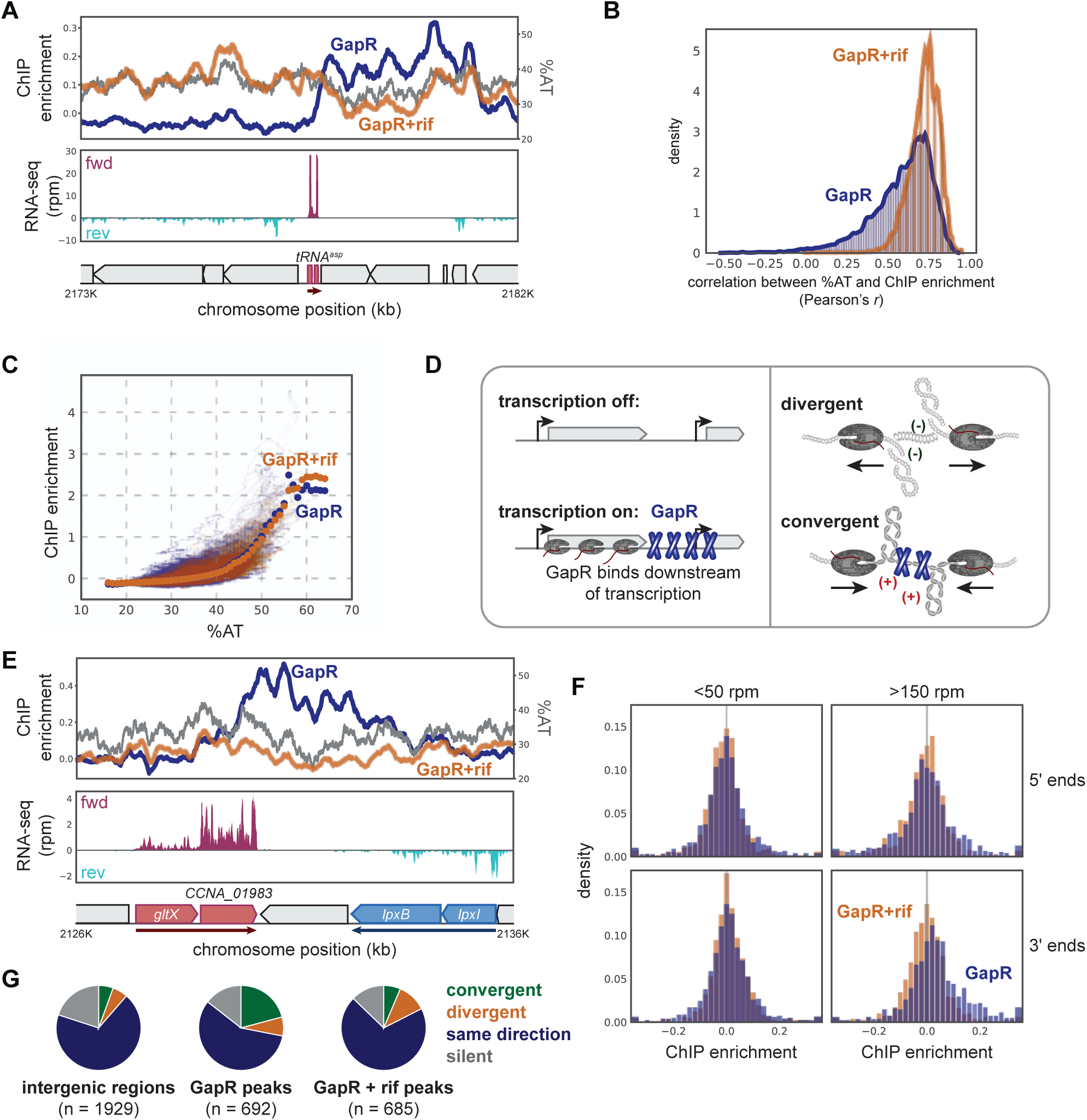
Transcriptional inhibition isolates AT-dependent GapR binding. (A) GapR -/+ rif ChIP at a highly expressed operon. Top: untreated (blue) and rif treated (orange) GapR profiles are plotted against average AT content (gray). Middle: transcription from the forward (red) and reverse (blue) strands. Bottom: positions of genes. (B) Pearson correlation comparing local AT content to GapR signal without (blue) or with rif treatment (orange). (C) GapR signal without (blue) or with rif treatment (orange) is plotted versus AT content across the genome. (D) GapR enrichment occurs at regions of known positive supercoiling including at the 3ʹ ends of highly transcribed genes and between active convergent genes. (E) GapR -/+ rif ChIP at a convergently-oriented locus as in (A). (F) GapR+rif ChIP is not enriched at the 3ʹ ends of highly transcribed operons. GapR accumulation was calculated at the 5ʹ and 3ʹ ends of operons separated by transcriptional activity. (G) GapR -/+ rif ChIP in different transcription orientations. Pie charts summarize the orientation of flanking genes for all intergenic regions (left), untreated ChIP peaks (middle) and rif-treated peaks (right).

The correlation between GapR signal strength and AT-content (17, 20) suggests that binding at AT-rich sites should increase upon rif treatment. To test this idea, we analyzed GapR enrichment genome wide. We find that GapR signal increases at AT-rich and decreases at AT-poor loci upon rif treatment (Fig. 8c, S2b). If GapR binding is mediated by AT content in the absence of +SC, we would predict that binding sites will be more AT-rich upon rif treatment. To test this prediction, we identified GapR and GapR+rif enriched and de-enriched sites generating 692 and 689 peaks and 1602 and 1905 valleys, respectively (see Methods).

GapR peaks are more AT-rich than valleys or a random sample of the genome, with peaks increasing in AT-richness upon rif treatment (Welch’s *t*-test: p < 10^-4^, Fig. S2c). In sum, our analyses show that in the absence of transcription-mediated +SC, AT-richness is the primary factor mediating GapR localization.

### AT-dependent binding occurs independently of +SC-dependent binding

To determine if the AT-rich GapR peaks we identified in rif-treated cells represent new GapR binding sites or a shift within existing sites, we assessed if GapR binding to +SC sites is altered upon treatment. We previously reported that +SC-dependent GapR binding occurs downstream (3′ ends) of highly active TUs and between convergently-oriented TUs (Fig. 8d) (20, 21). Indeed, at highly transcribed or convergently-oriented TUs, we find GapR signal is lost post rif-treatment (Fig. 8a, 8e), supporting the idea that GapR redistributes to new sites in the absence of +SC. We then quantified this behavior genome-wide. To measure GapR at TU ends, we classified TUs as highly active (>150 RPM) or poorly expressed (<50 RPM) and calculated GapR accumulation at the 5′ and 3′ end of the operon (see Methods). In untreated cells, GapR accumulates at the 3′ end of highly active TUs, but not at the 5′ ends or at the 5′ and 3′ of poorly expressed TUs as we previously noted (Fig. 8f) (20). Notably, upon rif treatment and loss of +SC, this 3′ accumulation is absent (Fig. 8f).

Next, to assess GapR at convergently-oriented TUs, we classified each GapR or GapR+rif peak to be between silent, same orientation, convergently-, or divergently-oriented TUs (see Methods). Convergent regions were highly enriched for GapR in wild type compared to rif-treated cells (Fisher’s exact test: p < 10^-11^, Fig. 8g). Rif treatment re-distributed GapR enrichment towards divergent regions (Fisher’s exact test: p < 10^-1^, Fig. 8g). Together, our analyses demonstrate that in untreated cells, GapR exhibits both AT-dependent and +SC-dependent binding, with GapR redistributing to AT-rich loci in the absence of transcription and +SC.

### GapR localization is largely dependent on +SC

To assess how often AT-dependent and +SC-dependent GapR binding occurred in untreated cells, we classified each GapR peak as mediated by AT only, +SC only, or by both AT and +SC. GapR peaks that decreased less than 5% upon rif treatment were considered mediated by AT only. Since the remaining peaks may have +SC only peaks and peaks mediated by both AT and +SC, we compared peaks to our GapR+rif dataset, classifying any that overlapped as mediated by both AT and +SC (see Methods, Fig. S3a-c).

To validate our classification method, we confirmed that: 1) AT only peaks are more AT-rich than +SC only peaks; 2) +SC only peaks are more enriched in convergent regions than AT only peaks; and 3) peaks mediated by both AT and +SC displayed intermediate phenotypes for both AT content and convergent region enrichment, indicating that these peaks are indeed mediated by both binding modes (Fig. S3d-e). Our approach yielded 107 AT only, 311 +SC only, and 274 peaks mediated by both AT and +SC (Fig. 9a). Our results show that both AT content and +SC mediate GapR binding, with the majority of GapR behavior dependent on +SC (85%) in untreated cells.

**Fig. 9.:**
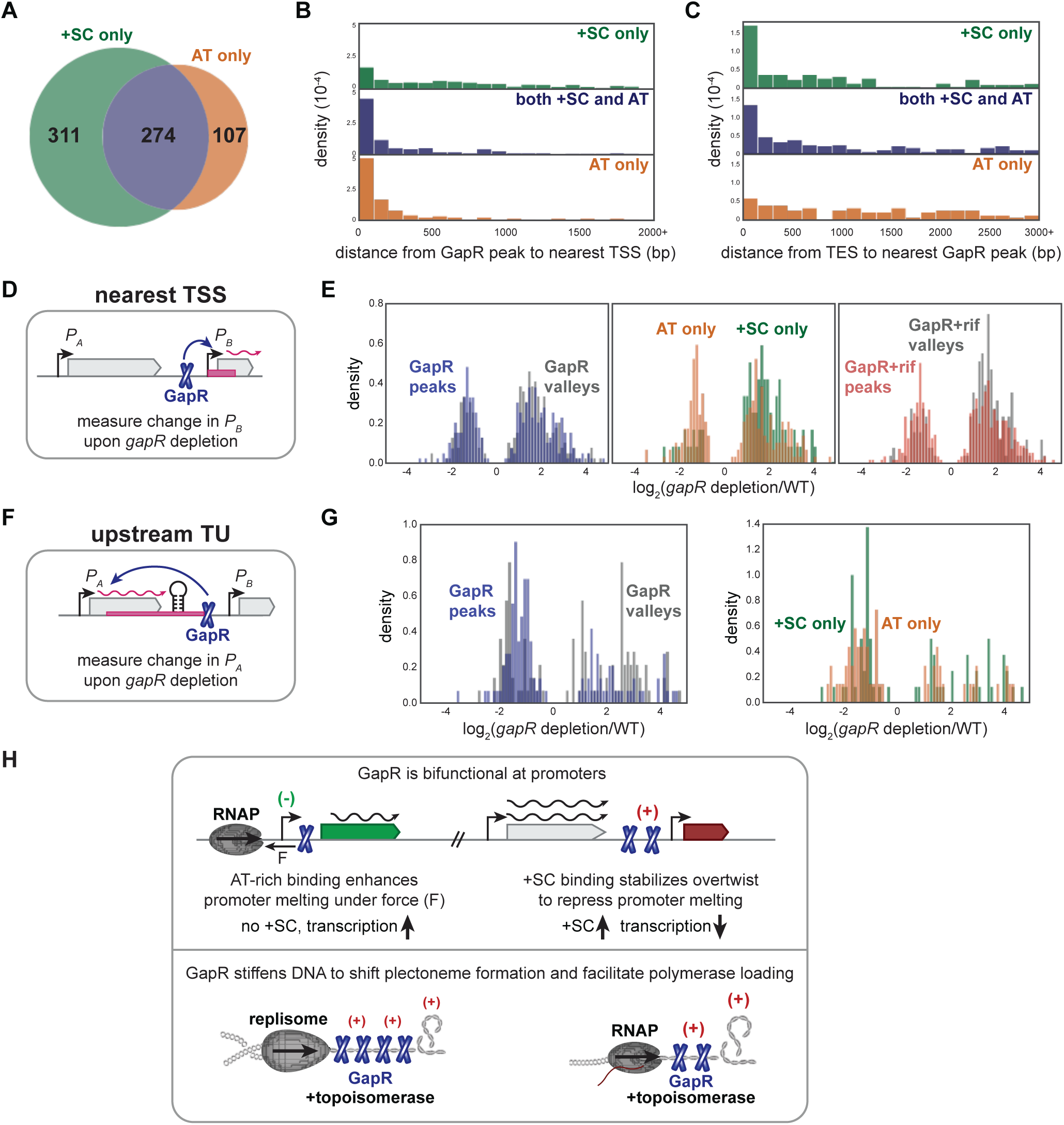
AT-rich and +SC-dependent binding promote RNAP progression but diferentially impact gene expression. (A) Venn diagram summarizing AT-rich and +SC-dependent GapR binding events in untreated cells. (B) Distance from GapR peaks to the nearest TSS. (C) Distance from all TES to the nearest GapR peak. (D) Analysis of GapR effect on transcriptional initiation. The nearest TSS to a GapR peak or valley is identified and the transcriptional activity of that TSS is calculated upon *gapR* depletion. (E) AT only and +SC only GapR binding have differential effects on transcription initiation. Histograms comparing log_2_-fold transcriptional change of the top 20% of GapR peaks or valleys upon *gapR* depletion. Untreated (left), AT only and +SC only GapR peaks (middle), and upon rif treatment (right). (F) Analysis of GapR effect on transcriptional elongation. The nearest upstream TSS to a GapR peak or valley is identified and the transcriptional activity between the upstream TSS and GapR peak or valley calculated upon *gapR* depletion. (G) GapR binding modestly stimulates transcription elongation. Histograms showing log_2_-fold transcriptional change upon *gapR* depletion. Left: GapR valleys versus peaks. Right: AT only versus +SC only events. (H) Models of GapR behavior influencing transcription initiation (top) and facilitating polymerase movement (bottom).

### AT-rich and +SC-dependent GapR binding both support RNAP progression but have differential effects on transcription

Other groups have proposed that GapR binds at promoters and other AT-rich regions to tune transcription (17–19, 22). However, our earlier work indicated no correlation between GapR binding and transcription (20). As we have separated AT only and +SC only GapR binding, we asked if either binding mode affects transcription by measuring the association of GapR binding with transcription start sites (TSS) or intrinsic transcription end sites (TES) (42, 43). Strikingly, AT only GapR peaks are significantly closer to TSS than +SC only peaks (Welch’s *t*-test: p < 10^-13^, Fig. 9b). In contrast, +SC only peaks are modestly, but not significantly closer to TES than AT only peaks (Welch’s *t*-test: p = 0.154, Fig. S4a). As +SC can accumulate in genomic regions other than at intrinsic terminators (e.g. Rho-dependent terminators), we performed a reciprocal analysis examining TES sites for GapR enrichment. Here, we observe a significant enrichment of +SC only peaks occurring near TES relative to AT only peaks (Welch’s *t*-test: p < 10^-39^, Fig. 9c). We also calculated the distance between TSS and GapR peaks, finding that while AT only binding is closely positioned to TSS, some +SC only peaks also occur close to TSS sites (Fig. S4b). As expected, mixed AT and +SC mediated peaks displayed an intermediate phenotype in all our analyses (Fig. 9b-c, S4a-b).

Because we identified GapR binding close to TSS, we wondered whether GapR would activate, repress, or have no or variable effects on promoter activity. To test this idea, we identified the nearest TSS to GapR peaks and asked if depletion of *gapR* would lead to a transcriptional change at this TSS (Fig. 9d). We used both a stringent cutoff and a more relaxed cutoff (5% and 20% of GapR enrichment, respectively) to capture all possible TSS. Strikingly, while GapR peaks are largely indistinguishable from valleys in their transcriptional effect, we find that AT only and +SC only peaks have modest, but significant and unique effects on transcription (Mann-Whitney test: p < 10^-4^, Fig. 9e, S4c-d). GapR appears to act as an activator at promoters adjacent to AT only peaks, while repressing promoters near +SC only peaks (Fig. 9e, S4c-d). Supporting this idea, we find that TSS associated with GapR+rif peaks also decrease in expression in the absence of *gapR* (Mann-Whitney test: p < 10^-2^, Fig. 9e, S4c-d). As each GapR binding mode has a unique effect on nearby promoter activity, we wondered if this would affect transcription surrounding the GapR binding site. Indeed, while GapR peaks and valleys are comparable in their transcriptional effect, AT only GapR peaks modestly, but significantly enhance local transcription while +SC only peaks repress it (Fig. S4e-f). These data show that GapR localization has two independent tuning effects on transcription: AT-dependent binding occurs near TSS sites to enhance transcription, while +SC-dependent binding occurs near TES sites and represses nearby promoter activity.

We previously proposed that GapR binds to the overtwisted DNA in front of the replisome and RNAP to stimulate topoisomerase activity and support efficient polymerase progression (20). Since we observed each binding mode to have a differential effect on initiation, we wondered if either binding mode supports RNAP elongation as described in Fig. 9f. We find that GapR modestly, but significantly supports transcription elongation, regardless of the binding mode (Fig. 9g). Taken together with our biophysical studies, we conclude that GapR is a unique NAP that binds both AT-rich and +SC DNA, stiffening DNA and promoting its melting, allowing GapR to regulate transcription initiation and promote polymerase progression.

## Discussion

α-proteobacterial GapR is an essential and pleiotropic NAP that promotes replication by regulating +SC. Previous studies showed GapR may bind to both AT-rich and +SC DNA, leading to conflicting models of GapR function as either a transcription factor or a topological sensor (17–21). Here, we validate with MT and BLI that GapR indeed has affinity for both types of DNA. Once bound, GapR stiffens DNA, increases its bending persistence length, and discourages plectoneme formation. By analyzing published ChIP-seq and RNA-seq datasets (20), we discriminate between these binding modes, revealing that each mode directs GapR to distinct loci with opposing effects on transcription initiation. Further, we suggest that both forms of GapR binding promotes replisome and RNAP elongation by holding the DNA in a rigid overtwisted conformation. Together, we propose that GapR has a primary role in promoting polymerase elongation and a secondary role in tuning transcription initiation.

### GapR stiffens DNA and is extremely stable on DNA

Under physiological levels of salt and force, we observed GapR binding to shift the relaxed DNA linking number (ΔLk_0_) and increase DNA stiffness (as reflected by the bending and twist persistence lengths, Figs. 2-3) in a concentration dependent manner. At slightly higher forces (0.5 pN), GapR facilitated double strand melting upon -SC, evident from an asymmetric extension versus linking number curve. GapR binding also altered the force-extension behavior of dsDNA, increasing its bending persistence length and decreasing its contour length. GapR also enhances the twist persistence length without markedly affecting pre-buckling curvature, while modestly increasing the energy required for plectoneme formation (Fig. 4).

Next, our kinetic experiments indicated that once GapR-DNA complexes were highly stable to washout, with little or no dissociation of protein observed over hours (Fig. 6). This is in accord with observations of similar stabilities of other protein-DNA complexes (12, 39). We found that partial dissociation could be driven using competing DNA fragments, showing that GapR displays “facilitated dissociation” behavior. Together, our data suggest that GapR binding limits local DNA bending and plectoneme formation. For GapR to be turned over *in vivo* it is likely that competitor factors such as newly synthesized DNA or active processes such as a moving replication fork are required.

### GapR stabilizes a conformation of DNA that promotes polymerase progression

GapR constrains DNA in a rod-like state that promotes duplex DNA melting and genes upstream of GapR binding sites are repressed in the absence of GapR. These properties lead us to propose that GapR may support replication and transcription directly by stiffening DNA to maintain a linearized template and facilitate its loading into these polymerases (Fig. 9h). Such behavior aligns with the suggestion by Arias-Cartin that GapR tracks with replication forks (19) and modulates DNA topology during replication or RNAP elongation. As the buckling point for plectonemes remains unchanged, this stiffening mechanism allows GapR to shift plectoneme formation away from the polymerases, enabling topoisomerases to access +SC ahead of these molecular machines.

We note that DNA stiffening by GapR could also contribute to the formation of local topological domains within a chromosome. These mechanical changes are reminiscent of the activities of other NAPs, such as H-NS and IHF, which also modify DNA stiffness to promote large-scale chromosomal organization. However, unlike H-NS, which represses transcription at AT-rich loci, GapR reinforces DNA organization without strictly repressing transcription (see below).

### GapR has affinity for AT-rich DNA and +SC DNA

Consistent with previous reports (17, 19, 20, 22, 23), our BLI experiments showed that GapR displayed preferential binding to AT-rich DNA, as indicated by lower *K_d_* and Hill coefficients, compared to GC-rich sequences (Fig. 5). However, our MT also indicated preferential binding to +SC DNA as we previously proposed (20, 21). At a DNA pulling force of 0.25 pN, GapR binding increases superhelical density by a maximum of 1.5%, consistent with protein-induced overtwisting. Binding in MT follows a Hill-like behavior, yielding an apparent K_d_ of 475 nM and a Hill coefficient of 3.25 (based on the shift in linking number), suggesting high cooperativity during GapR tetramer assembly, possibly without the more extensive filamentation that has been proposed)(24). Importantly, GapR exhibited enhanced affinity for overtwisted DNA (Fig. 7a), suggesting a feedback loop whereby +SC accumulation reinforces binding and potentiates additional DNA unwinding by replication and transcription complexes.

Our computational analysis of *in vivo* GapR binding validates and distinguishes between both binding modes. Most GapR binding sites are dependent on +SC only (45%), with a minority mediated AT-content only (17%), and the remainder (38%) mediated by both factors (Fig. 9a). Strikingly, when transcription and its generated supercoils are eliminated during rif treatment, GapR largely adopts the AT only binding mode. For events mediated by both +SC and AT-richness, the intensity of the AT-rich peak does not substantially decrease upon transcriptional inhibition, suggesting that both binding modes are independent. Our analyses thus provide an explanation for why prior ChIP-seq profiles differed in their ability to capture GapR behavior, as GapR will relocate and partition to AT-rich sites dynamically in response to transcription. We propose that AT-rich binding may represent a protein reservoir to correctly position GapR at sites where its activity will be needed as +SC occurs.

### GapR is a unique NAP that enhances and represses transcription

NAPs are known to manipulate DNA in ways that promote nucleoid organization and affect transcription. For example, NAPs like HU bend DNA to alter global transcription via incompletely described mechanisms. We show here that GapR functions differently from existing NAPs, with GapR binding tuning promoter activity depending on the genomic context and binding mechanism (sequence-vs topology-specific).

When GapR is bound at AT-rich, low +SC promoters, binding enhances transcription, as was previously suggested (17). We propose this occurs due to GapR binding (with concomitant overtwisting) inducing *undertwisting* of nearby DNA (as in the MT experiments, Fig. 4d), facilitating promoter melting by RNAP (Fig. 9h). This proposal is based on two pieces of evidence. First, the majority of AT-dependent GapR binding occurs <100 bp of TSS sites, placing it at an appropriate distance to mediate transcription initiation. Second, we show that GapR promotes dsDNA melting under relatively low tension (0.5 pN) in the presence of -SC. We hypothesize that translocating RNAP generates the force required to aid GapR in duplex melting.

Next, we find that AT-poor, +SC binding events also co-occur with promoters, with GapR binding at these sites repressing transcription. We propose that GapR traps local overtwist to prevent +SC from diffusing away, thus impeding promoter melting (Fig. 9h). Supporting this idea, we find that +SC-dependent binding occurs at greater distances (>100 bp) from TSS sites indicating transcriptional repression does not simply occur via promoter occlusion. These differential functions at AT-rich and +SC-dependent binding sites likely explain the conflicting literature as to whether GapR is a transcriptional regulator or a topological sensor. Our findings suggest GapR is both.

In sum, our work shows that GapR is a bifunctional NAP capable of both AT-rich and +SC-dependent binding. Both binding modes stiffen DNA and discourage plectoneme formation, allowing GapR to promote replisome and RNAP progression and modulate transcription initiation. These activities make GapR unique among NAPs (11). First, while GapR does recognize DNA sequence-specifically, GapR binding does not occlude promoters or recruit the transcription apparatus, meaning it is not a transcription factor or silencer.

Second, while GapR binds to a “bent” DNA shape, this binding does not loop DNA and cannot activate transcription directly, as DNA melting also requires force. Lastly, as GapR levels are limited (∼2000 copies/cell), its bending activity is insufficient to mediate global chromosome organization like HU (∼70, 0000 copies/cell) (20, 44). As GapR is restricted to the α-proteobacteria, our results raise the question of how widespread a “GapR-like” strategy for transcription and replication enhancement might be. We propose that other NAPs, e.g. IHF or Fis, may recognize both DNA sequence and topology, leading to context-dependent outcomes that could explain their diverse functions (2). Furthermore, as recognizing and responding to +SC is a universal cellular problem, we hypothesize that other NAPs may similarly recognize +SC to stiffen DNA and support transcription and replication progression.

## Acknowledgements

This work was supported by NIH grants R01-GM105847 (to J.F.M.), R35-GM154727 and R00-GM134153 (to M.S.G.), T32-GM136534 and NSF GRFP 22-614 (to L.M.K.), T32-GM008382 (to X.W.) and by subcontract to the University of Massachusetts under NIH grant UM1-HG011536 (Center for 3D Structure and Physics of the Genome, 4DN Consortium, to J.F.M.). We thank M. Schumacher and M. LeRoux for their comments on this manuscript.

## Code availability

Python scripts for the GapR computational analysis are available as a GitHub repository at https://github.com/msguo11/GapR_AT-SC-calling.

## Author contributions

X.W.: Conceptualization, Data curation, Formal analysis, Funding acquisition, Investigation, Methodology, Resources, Validation, Visualization, Writing – original draft, Writing – review & editing.

L.M.K.: Conceptualization, Data curation, Formal analysis, Funding acquisition, Investigation, Methodology, Software, Validation, Visualization, Writing – original draft, Writing – review & editing.

R.K.: Conceptualization, Data curation, Formal analysis, Investigation, Methodology, Validation.

J.F.M. and M.S.G.: Conceptualization, Data curation, Funding acquisition, Methodology, Project administration, Resources, Software, Supervision, Validation, Visualization, Writing – original draft, Writing – review & editing.

## Methods

### DNA substrate preparation

The primary DNA substrate for tethering was a linear, ∼11.4 kb double-stranded DNA fragment labeled at each end. This DNA tether was generated from plasmid pNG1175 (9702 bp, a derivative of pFOS-1). The plasmid was linearized by digestion at unique SpeI and ApaI sites, yielding a 9438 bp fragment. Each end was then ligated to a ∼0.8 kb PCR product containing either biotinylated or digoxigenin-labeled nucleotides with SpeI- or ApaI-compatible overhangs. The resulting construct was a ∼11.4 kb linear DNA labeled with biotin at one end and digoxigenin at the other. This labeled DNA was used as the tether in single-molecule experiments.

### Buffers and solution conditions

All experiments were performed in 20 mM Tris buffer containing 100 mM NaCl. All buffers were adjusted to pH 7.5 and filtered before use. Sheared herring sperm DNA (HS-DNA, Promega) was used as a nonspecific competitor in kinetic experiments. Giuntoli et al. reported fragment sizes averaging ∼750 bp, ranging from ∼100 bp to ∼3 kb. Short specific DNA oligonucleotides were also occasionally tested as competitors.

### Flow cell assembly

Single-molecule experiments were conducted in custom flow cells assembled for each trial. A thin strip of parafilm, cut slightly smaller than the glass slide, was slit longitudinally to create a narrow flow channel. An elliptical opening was created at the midpoint of the channel. The resulting flow cell volume was approximately 30-40 μL. This parafilm layer was placed onto a glass slide, and a coverslip was positioned on top, ensuring both ends of the slit remained exposed. Liquid samples were introduced by capillary action.

### Surface functionalization and passivation

Flow cell surfaces were prepared by antibody coupling following standard protocols. The top surface of each flow cell was functionalized with anti-digoxigenin antibodies (Roche Diagnostics) diluted in PBS. The assembled flow cells were stored in a humid chamber and incubated overnight to capture digoxigenin-labeled DNA ends. The flow cells were then passivated for 2 hours with 1% BSA in PBS to prevent nonspecific adsorption of proteins or DNA to chamber walls. Excess BSA was subsequently rinsed away using 100 mM NaCl Tris buffer.

### DNA attachment to flow cell and to paramagnetic beads

To form DNA tethers, the prepared ∼11.4 kb biotin/digoxigenin-labeled DNA was first mixed with streptavidin-coated paramagnetic beads (2.7 μm diameter, Dynabeads M-270, Invitrogen) in Tris buffer. Biotinylated DNA ends rapidly bound to streptavidin on the bead surfaces. After incubation for 15 minutes, the DNA–bead complexes were introduced into the passivated flow cell. The digoxigenin-labeled free end of each DNA molecule readily attached to the anti-digoxigenin-coated coverslip, anchoring one end of the DNA to the surface and tethering the other end to a magnetic bead. Excess unbound beads and DNA were gently washed out with Tris buffer, and the flow cell was refilled with fresh buffer.

Successful tether formation was confirmed visually by observing beads that responded magnetically yet remained bound near the surface via a single DNA molecule.

### Vertical magnetic tweezers experiments

Once DNA tethers were established within the flow cell, the chamber was mounted on a vertical magnetic tweezer platform. The experimental setup consisted of a microscope equipped with a high-numerical-aperture objective (100×, NA 1.3 NA, Olympus), positioned on a piezoelectric stage (MIPOS 100 sge) to allow precise focal adjustments above the flow cell. Below the flow cell, a pair of permanent neodymium magnets was placed using a motorized stepper-motor-driven translator, aligning along the vertical axis of the microscope length. Tension on the DNA tether (stretching force) could be systematically applied and varied to the magnetic bead–DNA tether system by adjusting the magnet-to-flow cell distance.

Real-time tracking of the vertical positions of tethered beads is achieved by a camera attaching to the objective lens and image analysis algorithms. Polystyrene beads adhering to the glass surface provide fixed reference points for maintaining focus and correcting for mechanical drift. The bead height, corresponding to DNA extension, is determined using an auto focus algorithm that analyzed diffraction ring pattern, evaluating focus sharpness, and also by a separate algorithm that uses the degree of focus of the beads to determine their distance from the glass, both implemented by Labview (National Instruments, Austin TX) programs. Both algorithms can quantify the bead’s axial position relative to the focal plane with sub-micron precision.

Force calibration is achieved by an established method (12, 45), utilizing thermal fluctuations of the bead position. Thermal fluctuations in the bead’s position were recorded and converted to force values to calculate force value. The proper bead-DNA attachment is examined using the known force-extension relation for DNA molecule (35). Additionally, supercoiling capability is verified by observing the relation between rotations of the magnets on DNA extension, eliminating possible damaged, nicked DNA molecule.

### Supercoiling experiments

After confirming that the tether corresponds to a single DNA molecule, a typical experiment begins at a fixed force of 0.9 pN, with the linking number (ΔLk) systematically scanned from –40 to +40 in steps of 5 turns in experimental buffer (100 mM NaCl in Tris, without protein). Each step is approximately 10-15 seconds. Subsequently, two additional scans are performed: first from –35 to +35 at 0.5 pN (step size of 5), and then from –30 to +30 at 0.25 pN (step size of 2). During these measurements, the DNA tether extension (z) and corresponding linking number (ΔLk) are continuously monitored and recorded using LabVIEW software. Following this, 150 *μL* GapR protein solution (4-6 times the volume of flow cell) is introduced into the flow cell in GapR buffer (Tris buffer containing 100 mM NaCl). Finally, the DNA tether extension is measured again at each linking number for the same three forces (0.9 pN, 0.5 pN, and 0.25 pN) after about 5-10 minutes for buffer exchange and protein binding.

### Force-extension experiments

A slightly modified procedure is used to study the effect of the GapR protein on DNA contour length and persistence length. After confirming that the tether corresponds to a single DNA molecule, the experiment is initiated at an initial magnet position corresponding to ∼5.5 pN pulling force followed by scanning the vertical position of the neodymium magnet, which produces a force scan ranging from 0.1 to 5.5 pN. The extension of the DNA molecule and the magnet’s position are then measured and recorded. To ensure that these changes are not due to supercoiling induced by the GapR protein, a nicked DNA molecule (featuring a break in one strand) is used, thereby preventing melting or supercoiling from affecting the DNA’s extension. Subsequently, the 150 *μL* GapR protein is introduced into the flow cell in GapR buffer (a Tris buffer containing 100 mM NaCl), and the DNA tether extension is measured again at the same magnetic position.

### Kinetic Magnetic Tweezers Experiments

In “kinetic” MT experiments, following binding of GapR to DNA, an extension-linking number curve was measured. After that, the protein solution was subsequently exchanged with 200 *μL* protein-free buffer, and the length of the tethered DNA was re-measured at the given linking number under tensions of 0.25 pN, 0.5 pN, and 0.9 pN after 90 minutes.

### Comparing protein binding at different linking numbers

To test whether the GapR protein exhibits a binding preference for +SC DNA, the DNA tether was first exposed to 400 nM GapR protein at a zero-linking-number (torsionally relaxed) under a pulling force of 1 pN for 10 minutes. Following this incubation, the protein solution was removed by flushing the flow cell with 150 *μL* Tris buffer—using a volume approximately five to six times that of the flow cell. After 10 minutes in the GapR-free Tris solution, an extension–linking number measurement was taken at pulling forces of 0.9 pN, 0.5 pN, and 0.25 pN. Next, the linking number was adjusted to +15 while maintaining a pulling force of 1 pN. The same GapR solution, in a volume approximately five times that of the flow cell, was then introduced to replace the binding buffer for about 10 minutes. Finally, the GapR solution was replaced once again with GapR-free Tris buffer, and an extension– linking number measurement was recorded at 0.9 pN, 0.5 pN, and 0.25 pN after 10 minutes.

### BLI experiments

Magnetic tweezer experiments were supplanted with direct measurements of protein binding using biolayer interferometry (BLI). These experiments were conducted using an Octet N1 instrument using its dedicated software, with all buffers and biosensors equilibrated to room temperature. A 50-base-pair biotin labeled double-stranded DNA was used to test binding affinities for sequences containing 20%, 50%, and 80% G-C. The DNA was diluted by 100 times and loaded onto streptavidin coated sensor in a cuvette. The reference buffer contained 20% Tris, 100 mM NaCl, and 0.1% BSA, with the BSA added to prevent GapR from binding to the streptavidin sensor.

The sensors were hydrated in the reference buffer for 10 minutes before the experiments. The hydrated sensors were then immersed in reference buffer for 60 seconds to establish a stable baseline, monitored in real time through the interferometric signal.

Following this, the sensors were transferred to the DNA solution for 120 seconds to immobilize the dsDNA fragments onto the sensor surface. After DNA loading, the sensors were returned to the reference buffer for an additional 60 seconds to wash away unbound dsDNA molecules, stabilizing the signal for subsequent steps.

For the association phase, the sensors were moved to wells containing the GapR, where binding interactions were monitored over 3 -12 minutes as GapR proteins bind to the dsDNA. Finally, to measure dissociation kinetics, the sensors were re-immersed in reference buffer or buffer containing herring sperm DNA for 10 minutes, allowing real-time observation of ligand-analyte complex separation. Throughout the process, the instrument’s software continuously tracked signal shifts, converting them into binding curves for kinetic analysis.

## Data analysis

The experimental data obtained from magnetic tweezers were analyzed to characterize the mechanical properties of DNA-protein complexes. Specifically, we investigated the relationships between extension, linking number, and force under defined buffer conditions (100 mM NaCl, Tris, pH 7.4), employing both magnetic tweezer setups.

### Determining persistence length and contour length from force-extension relation

Magnetic tweezers provided data for force-extension relationships under conditions of varying tension. To interpret these data, force-extension curves were fitted to the twistable worm-like chain (TWLC) model, 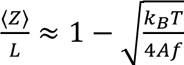, which effectively describes the mechanical behavior of semi-flexible polymers like DNA. By fitting the experimental force-extension data to this simplified TWLC model, we extracted mechanical parameters, including the DNA tether’s persistence length and contour length, allowing quantification of the DNA bending stiffness changes upon protein binding. To ensure the measured mechanical property changes were solely due to protein interaction and not supercoiling effects, nicked DNA substrates (containing a single-strand break) were used to prevent torsional stress accumulation.

### Determining DNA-Protein Binding Relation from Extension-Linking Number Curve

Data from magnetic tweezer experiments were analyzed to study the supercoiling and buckling characteristics of both DNA and DNA–protein complexes. In these experiments, the relationship between the extension of the DNA tether and its linking number in the stretched (pre-buckling) state follows a quadratic behavior (26, 34). By fitting a quadratic function to the top 15–20% of data points—where the reduction in DNA length is not yet absorbed into plectonemes—the relaxed linking number is determined. Changes in the peaks of this quadratic fit can be used to quantify the binding of proteins to double-stranded DNA, as protein binding can modify the relaxed linking number and, consequently, the shifting the extension–linking number relationship.

### Determining Twist Persistence Length of DNA-Protein Complex from Extension-Linking Number Curve

The curvature of the extension–linking number curve not only helps establish the relaxed linking number but also provides a means to extract the twist persistence length of the DNA–protein complex. According to prior work (26, 34), the average extension 〈Z〉 relative to the contour length L of the DNA tether is given by

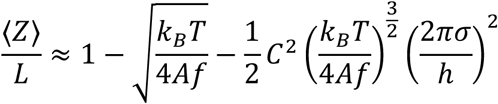

By taking the second derivative of the extension with respect to the linking number, one can calculate the curvature, K, of the extension versus linking number curve pre-buckling. This derivative can be algebraically transformed to solve for the twist persistence length, yielding

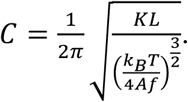

*C* quantifies the stiffness of the DNA–protein complex with respect to twist deformations, and any changes from the bare DNA behavior can be attributed to protein binding.

### Determining Plectonemic Persistence Length of DNA-Protein Complex from Extension-Linking Number Curve

The formation of plectonemes in the buckled regime of the extension–linking number curve offers another window into the mechanical properties of the DNA–protein complex. In the model described by Marko (26). the slope of the extension-linking number curve in the coexistence region (where both buckled and stretched states coexist) is linear. This relationship is expressed as *z* = *x_s_z_ext_*. where *x_s_* is the fraction of DNA remaining in the extended state and *x_s_z_ext_* is the extension at the buckling transition. Experimentally, *z_ext_* is obtained by identifying the intersection between the quadratic fit from the pre-buckling (top 15–20% of data points) region and the linear fit corresponding to the post-buckling regime. By defining the constant slope as 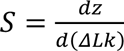, the plectonemic persistence length (p) can be calculated from 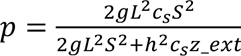,

where *g* and *c_s_* are model-specific constants related to the DNA’s mechanical properties. Determining both the twist and plectonemic persistence lengths allows one to theoretically predict the buckling point through torque equilibrium calculations. The torque τ in stretched DNA is given by 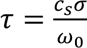, while in the coexistence region it takes the form 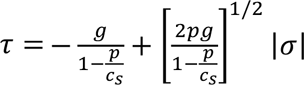 Marko (26). Accordingly, the linking number density at the buckling point can be expressed as 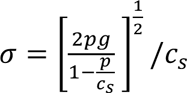.

Separating GapR binding modes.

All ChIP-seq (untreated, rif-treated) datasets were from GSE100657 (GSM2690549, GSM2690550, and GSM2690551) (20). ChIP-seq data was smoothed, normalized to reads per million, and mapped to the *Caulobacter crescentus* NA1000 reference genome (NC011916.1) as described previously (20). ChIP traces were then normalized by subtracting the GapR untagged control from the GapR-3xFLAG -/+ tagged datasets.

To identify GapR enriched loci, we isolated the top 5% or 20% of normalized GapR signal. Peaks began when the signal rose above the appropriate threshold and ended once the signal fell below threshold. Sites of de-enrichment were defined as regions that were in the bottom 5% or 20% of normalized GapR signal. To split the binding events into AT-dependent or +SC-dependent events, the location of the maximum ChIP signal in each GapR enrichment window within the untreated ChIP dataset was recorded. The peak was considered +SC-dependent if the maximum ChIP signal decreased by less than 5% upon rif treatment. The peak was AT-dependent if the enrichment window in which it was found also contained the location of a peak (max signal within an enrichment window) from the rif-treated dataset using the same threshold. Enrichment windows that satisfied both conditions were classified as both AT and SC-dependent.

### Measuring correlation with AT-richness

AT content at each base pair was calculated with a centered 300 bp sliding window. To identify correlations between AT content and GapR binding, AT content was plotted on linear and log_2_-scale versus normalized GapR ChIP at each position. The repetitive rDNA loci were removed from this analysis. For S2c and S3d, the average AT content of the 300 bp (flanking 150bp each side) surrounding the maximum GapR signal and displayed with box-and-whisker plots displaying the median and interquartile range (25th-75th percentile).

Pearson correlation was calculated between the local AT content (as above) and either the untreated or rif-treated ChIP signals across a 10 kb sliding window approach. Correlations are calculated only for positions where a complete 10kb window exists. Distributions were visualized using density-normalized histograms with overlaid kernel density estimates for both untreated and rif-treated datasets.

### Transcriptional orientation analysis

RNA-seq datasets were from GSE100657 (GSM2690552, GSM2690553, and GSM2690554) were analyzed as described previously (20). For the transcription unit analysis in Fig. 8g and S3e, intergenic regions were classified based on the relative orientation of the flanking transcription units (TU) (convergent, divergent, same direction, or silent) as was done previously (21). TU boundaries were defined as a continuous genomic region that is expected to be transcribed as a single RNA molecule, corresponding to a multi-gene operon or a single gene unit with strand specific start and end coordinates. Operon annotations to direct these boundaries were sourced from MicrobesOnline (46) . The average summed RNA-seq signal across the intergenic region and its 5000bp flanking regions must be below 0.1 to be considered transcriptionally silent. For same direction, divergent, or convergent classification, a 2000bp flanking region and strand specificity was used. Divergent regions are where the left side transcription is reverse-strand dominated and the right strand is forward-strand dominated or inversely. This logic was extended to define convergent and same direction-oriented TUs.

3′ and 5′ GapR accumulation as seen in Fig. 8f were quantified by collecting strand-specific 1kb sliding window averages of normalized ChIP flanking the immediate upstream and downstream binding of both the start and end sites of each TU. For each TU boundary, the difference between flanking windows were plotted in overlaid histograms for the 5′ and 3′ ends. TUs were stratified into highly expressed (>150 RPKM) and lowly expressed (<50 RPKM) units based on expression through the gene body. Distributions of the differences in TU accumulation were compared using independent two-sample *t*-tests comparing the high expression and low depression TUs for both the 5′ and 3′ accumulation.

### Measuring distance from TES and TSS

Transcription end sites (TES) and transcription start sites (TSS) were sourced from (42, 43). Distance from GapR to the nearest TES or TSS as seen in Fig. 9b and S4a are defined as distance of the maximum chip peak or alley and the nearest TSS or TES. For histograms in Fig. 9c and S4b, the distance from every TSS or TES to the nearest maximum GapR ChIP peak or valley was identified. For all histograms described, if the distance was larger than the graph axis limit, the value of the axis limit was recorded instead. For all analyses, statistical significance was calculated using Welch’s *t*-test.

### Measuring changes in transcriptional activity

Log_2_ fold change comparisons were done between wild-type log-phase growth in glucose compared to growth of a strain in which expression of *gapR* is repressed for 6 hours with glucose. For all analyses, statistical significance was calculated using the Mann-Whitney-Wilcoxon test.

For Fig. 9e and S4d, for each peak, the nearest TSS was identified the maximum log_2_- fold transcriptional change in the 500 bp downstream of the TSS was calculated. All duplicate TSS were removed so that plots and statistical tests represent unique, non-overlapping TSS sites. To measure GapR effect during transcription elongation, the nearest TES was found in the same manner as described for the nearest TSS, and an “elongation window” was defined as follows: the TSS that initiates transcription through the identified TES and the elongation window was defined starting 100bp downstream of the TSS to the position of maximum GapR ChIP signal. The maximum log_2_-fold transcriptional change within this window was recorded. Only windows where the GapR max ChIP signal was within 500bp of the predicted TES site were considered. Lastly, to measure log_2_-fold transcriptional changes under GapR peaks and valleys in Fig S4f, the largest log_2_-fold change was recorded within a 1kb window centered along the max ChIP peak or valley.

## Supplementary Materials

**Fig. S1.**
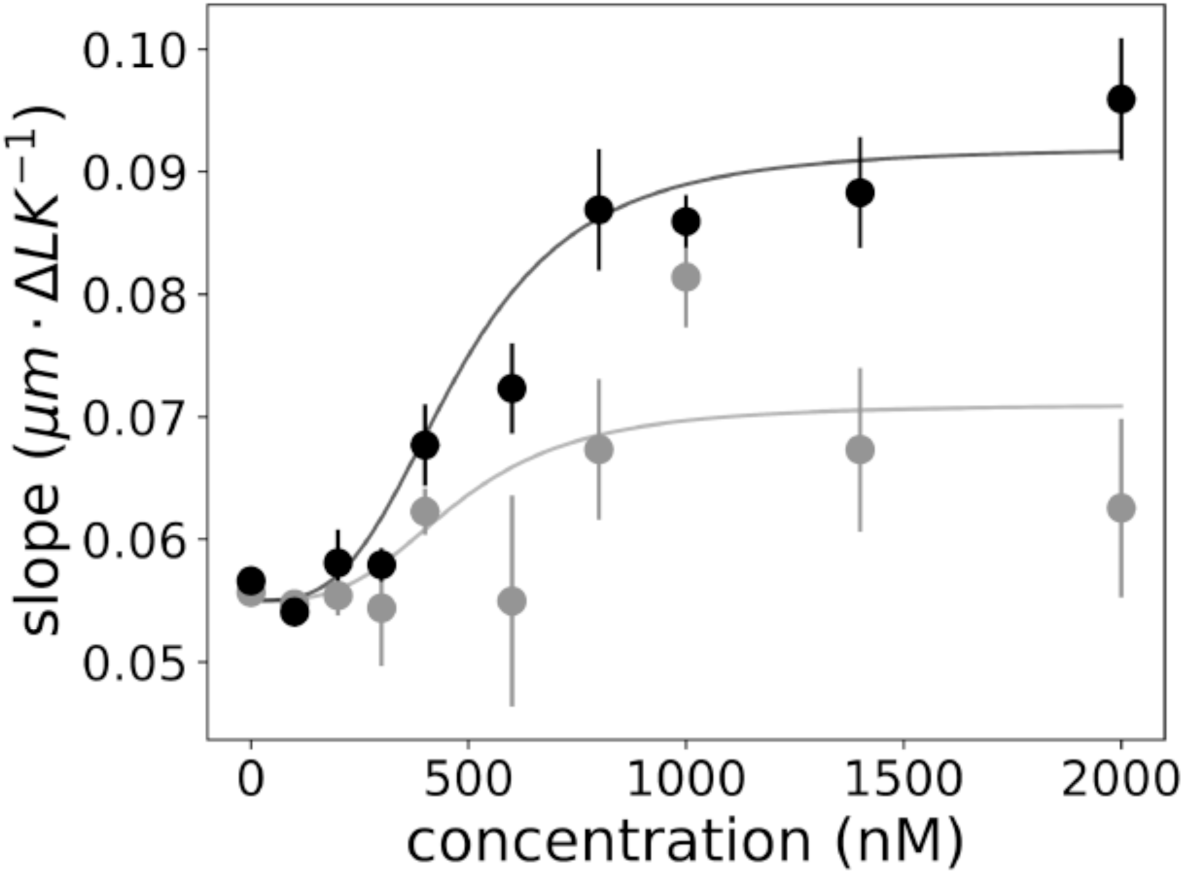
Slope of the extension-linking number curve at 0.25 pN within the coexistence region (post-buckling). This slope is utilized to calculate the plectonemic twist persistence length at 0.25 pN and the theoretical buckling point at 0.25 pN. Black: The average slope of the linking number-extension relation in the +SC plectonemic region. Gray: The average slope in the -SC plectonemic region. Solid lines indicate theoretical titration curves derived from Fig. 2b, employing the same Hill coefficient and dissociation constant (K_d_).

**Fig. S2.**
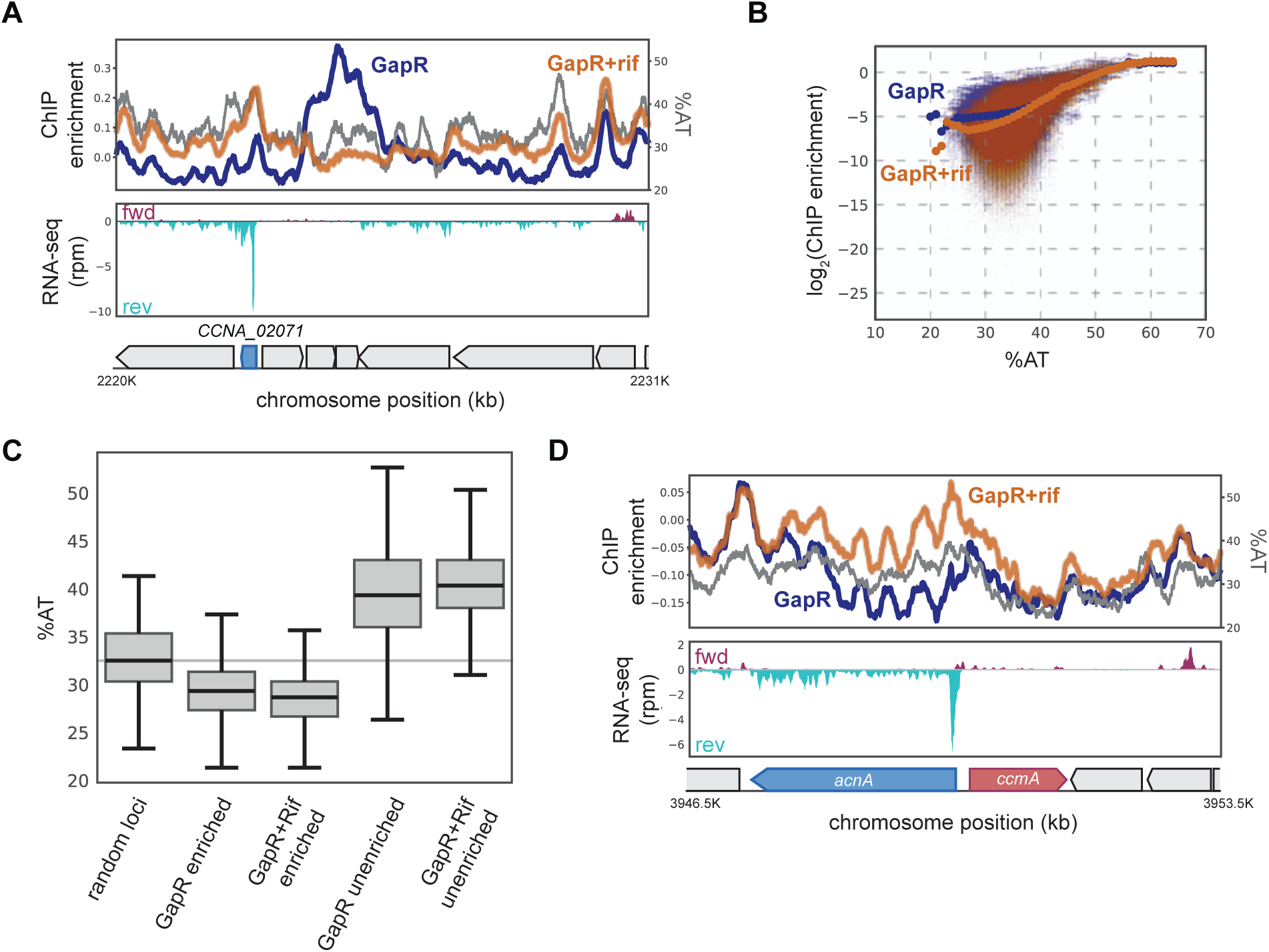
(A) GapR -/+ rif ChIP at a representative operon. Top: untreated (blue) and rif treated (orange) GapR profiles are plotted against average AT content (gray). Middle: transcription from the forward (red) and reverse (blue) strands. Bottom: positions of genes. (B) GapR signal without (blue) or with rif treatment (orange) is plotted versus AT content across the genome. (C) Box plots comparing average AT content within GapR peaks, GapR valleys, and the average of 100 random genomic loci. (D) GapR -/+ rif ChIP at a representative de-enriched locus as in (A).

**Fig. S3.**
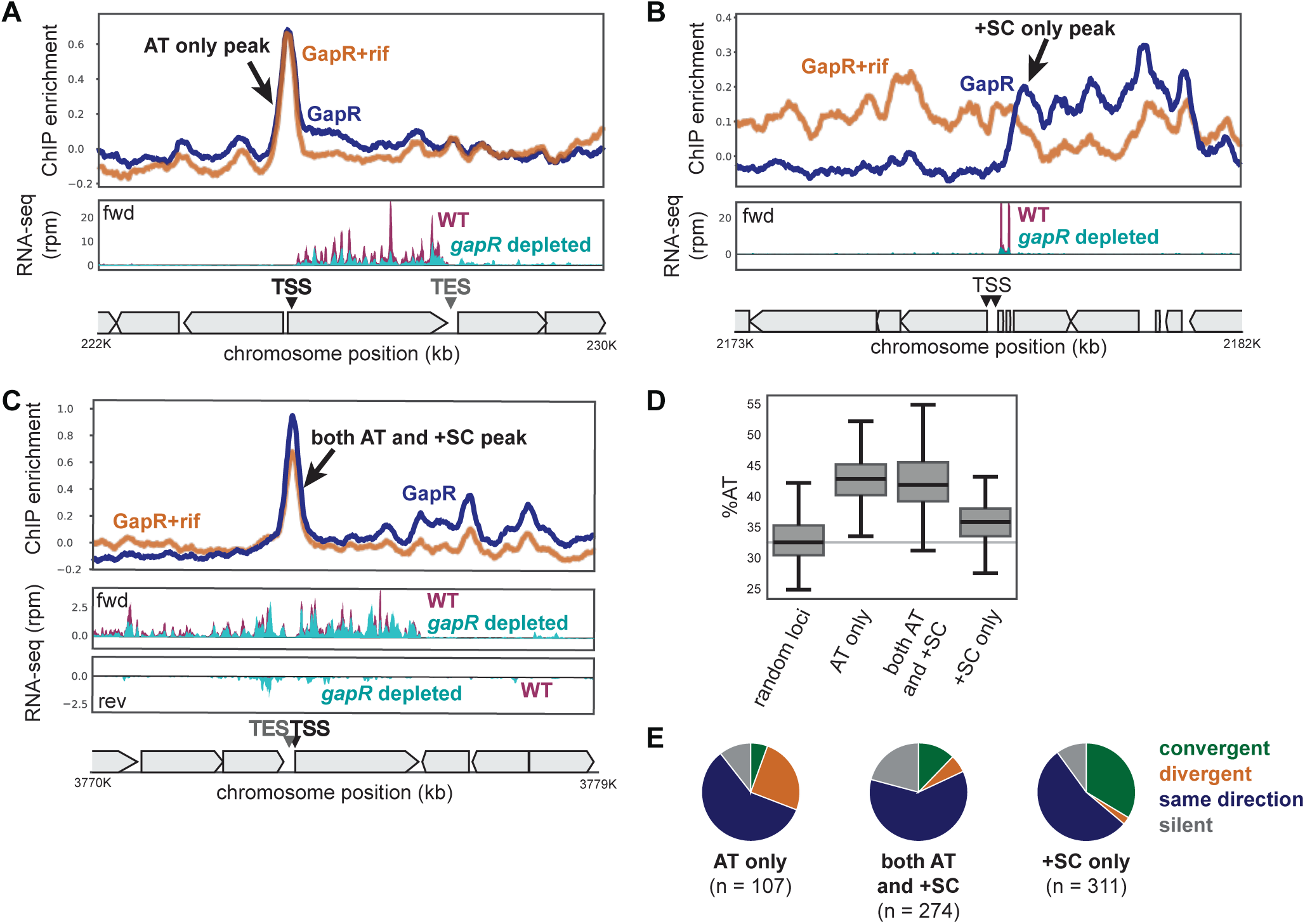
(A) GapR -/+ rif ChIP at an AT-rich binding event. Top: untreated (blue) and rif treated (orange) GapR profiles are plotted. Middle: transcription in wild-type and gapR depeleted cells. Bottom: TSS, TES, and positions of genes. (B) GapR binding at a +SC-dependent event as in (A). (C) GapR binding at an AT and +SC-dependent event as in (A). (D) Box plots comparing average AT content within GapR peaks and the average of 100 random genomic loci. (E) Pie charts summarizing the orientation of flanking genes for all AT only (left), both AT and +SC (middle), and +SC only peaks (right).

**Fig. S4.**
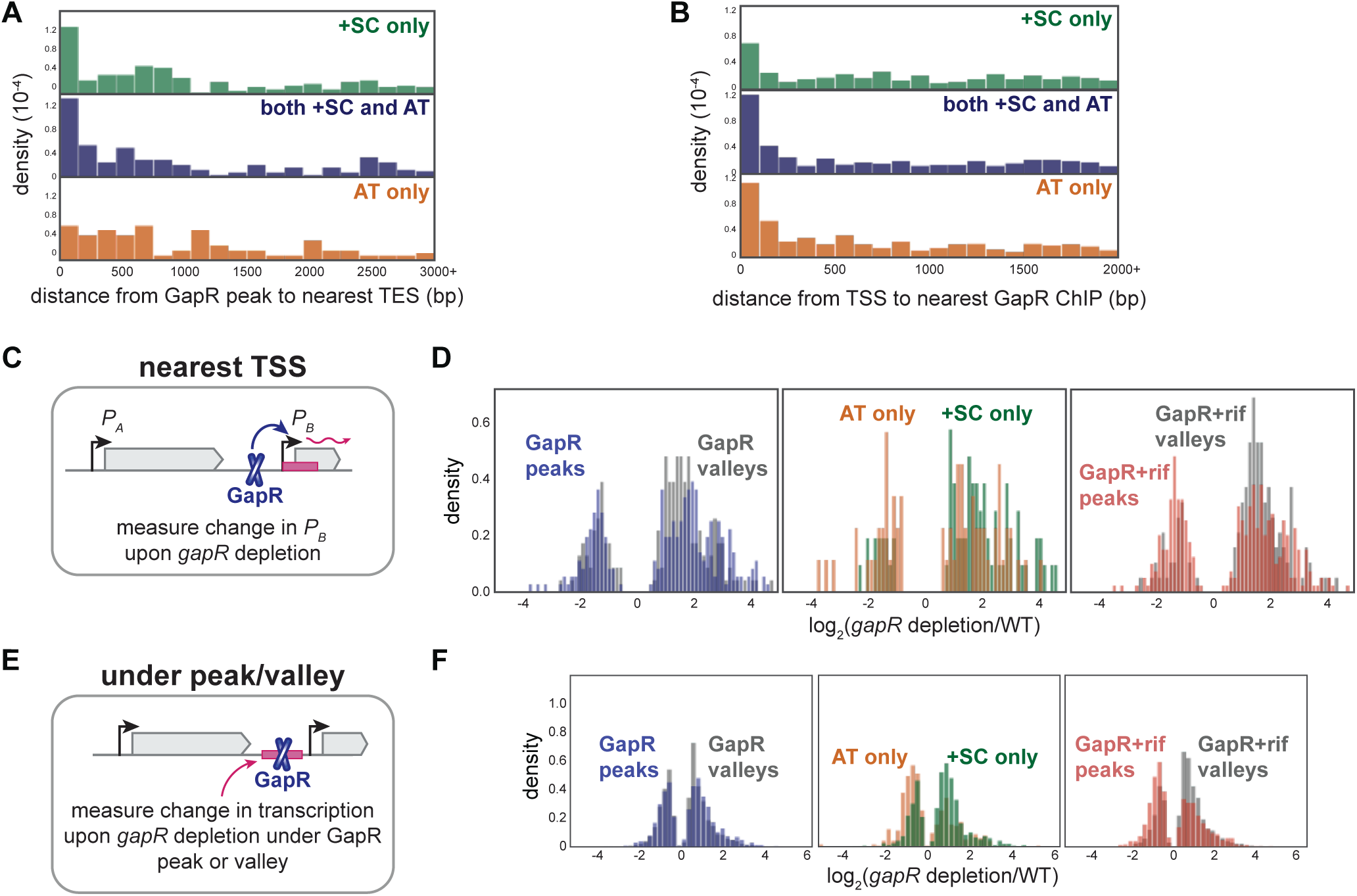
(A) Distance from GapR peak to nearest TES. (B) Distance from TSS to nearest GapR peak. (C) Analysis of GapR effect on transcriptional initiation. The nearest TSS to a GapR peak or valley is identified and the max transcriptional activity of that TSS is calculated upon *gapR* depletion. (D) Histograms as in Figure 9E comparing log_2_-fold transcriptional change of the top 5% of GapR peaks or valleys upon *gapR* depletion. Untreated (left), AT only and +SC only GapR peaks (middle), and upon rif treatment (right). (E) Analysis of GapR effect on local transcription. Max transcriptional activity upon gapR depletion in a 1 kb window centered at each GapR peak or valley is calculated. (F) Histograms showing the effect of GapR binding on transcription. Untreated (left), AT only and +SC only GapR peaks (middle), and upon rif treatment (right).

## References

1. Dame, R.T., Rashid, F.-Z.M. and Grainger, D.C. (2020) Chromosome organization in bacteria: mechanistic insights into genome structure and function. Nat. Rev. Genet., 21, 227–242.

2. Verma, S.C., Qian, Z. and Adhya, S.L. (2019) Architecture of the Escherichia coli nucleoid. Plos Genet, 15, e1008456.

3. Dillon, S.C. and Dorman, C.J. (2010) Bacterial nucleoid-associated proteins, nucleoid structure and gene expression. Nat Rev Microbiol, 8, 185–195.

4. Badrinarayanan, A., Le, T.B.K. and Laub, M.T. (2015) Bacterial Chromosome Organization and Segregation. Annu Rev Cell Dev Bi, 31, 171–199.

5. Hustmyer, C.M. and Landick, R. (2024) Bacterial chromatin proteins, transcription, and DNA topology: Inseparable partners in the control of gene expression. Mol. Microbiol., 122, 81–112.

6. Postow, L., Crisona, N.J., Peter, B.J., Hardy, C.D. and Cozzarelli, N.R. (2001) Topological challenges to DNA replication: Conformations at the fork. Proc National Acad Sci, 98, 8219–8226.

7. Wu, H.-Y., Shyy, S., Wang, J.C. and Liu, L.F. (1988) Transcription generates positively and negatively supercoiled domains in the template. Cell, 53, 433–440.

8. Liu, L.F. and Wang, J.C. (1987) Supercoiling of the DNA template during transcription. Proc National Acad Sci, 84, 7024–7027.

9. McKie, S.J., Neuman, K.C. and Maxwell, A. (2021) DNA topoisomerases: Advances in understanding of cellular roles and multi-protein complexes via structure-function analysis. BioEssays, 43, e2000286–e2000286.

10. Sutormin, D.A., Galivondzhyan, A.K., Polkhovskiy, A.V., Kamalyan, S.O., Severinov, K.V. and Dubiley, S.A. (2021) Diversity and Functions of Type II Topoisomerases. Acta Naturae, 13, 59–75.

11. Amemiya, H.M., Schroeder, J. and Freddolino, P.L. (2021) Nucleoid-associated proteins shape chromatin structure and transcriptional regulation across the bacterial kingdom. Biochem Soc Symp, 12, 182–218.

12. Skoko, D., Wong, B., Johnson, R.C. and Marko, J.F. (2004) Micromechanical Analysis of the Binding of DNA-Bending Proteins HMGB1, NHP6A, and HU Reveals Their Ability To Form Highly Stable DNA−Protein Complexes †. Biochemistry-us, 43, 13867–13874.

13. Noort, J. van, Verbrugge, S., Goosen, N., Dekker, C. and Dame, R.T. (2004) Dual architectural roles of HU: Formation of flexible hinges and rigid filaments. Proc. Natl. Acad. Sci., 101, 6969–6974.

14. Gulvady, R., Gao, Y., Kenney, L.J. and Yan, J. (2018) A single molecule analysis of H-NS uncouples DNA binding affinity from DNA specificity. Nucleic Acids Res., 46, 10216– 10224.

15. Schwab, S. and Dame, R.T. (2025) Identification, characterization and classification of prokaryotic nucleoid-associated proteins. Mol. Microbiol., 123, 206–217.

16. Prieto, A.I., Kahramanoglou, C., Ali, R.M., Fraser, G.M., Seshasayee, A.S.N. and Luscombe, N.M. (2012) Genomic analysis of DNA binding and gene regulation by homologous nucleoid-associated proteins IHF and HU in Escherichia coli K12. Nucleic Acids Res., 40, 3524–3537.

17. Ricci, D.P., Melfi, M.D., Lasker, K., Dill, D.L., McAdams, H.H. and Shapiro, L. (2016) Cell cycle progression in Caulobacter requires a nucleoid-associated protein with high AT sequence recognition. Proc National Acad Sci, 113, E5952–E5961.

18. Taylor, J.A., Panis, G., Viollier, P.H. and Marczynski, G.T. (2017) A novel nucleoid-associated protein coordinates chromosome replication and chromosome partition. Nucleic Acids Res, 45, gkx596-.

19. Arias-Cartin, R., Dobihal, G.S., Campos, M., Surovtsev, I.V., Parry, B. and Jacobs-Wagner, C. (2017) Replication fork passage drives asymmetric dynamics of a critical nucleoid-associated protein in Caulobacter. Embo J, 36, 301–318.

20. Guo, M.S., Haakonsen, D.L., Zeng, W., Schumacher, M.A. and Laub, M.T. (2018) A Bacterial Chromosome Structuring Protein Binds Overtwisted DNA to Stimulate Type II Topoisomerases and Enable DNA Replication. Cell, 175, 583–597.e23.

21. Guo, M.S., Kawamura, R., Littlehale, M.L., Marko, J.F. and Laub, M.T. (2021) High-resolution, genome-wide mapping of positive supercoiling in chromosomes. Elife, 10, e67236.

22. Huang, Q., Duan, B., Dong, X., Fan, S. and Xia, B. (2020) GapR binds DNA through dynamic opening of its tetrameric interface. Nucleic Acids Res., 48, gkaa644-.

23. Tarry, M.J., Harmel, C., Taylor, J.A., Marczynski, G.T. and Schmeing, T.M. (2019) Structures of GapR reveal a central channel which could accommodate B-DNA. Sci Rep-uk, 9, 16679.

24. Lourenço, R.F., Saurabh, S., Herrmann, J., Wakatsuki, S. and Shapiro, L. (2020) The Nucleoid-Associated Protein GapR Uses Conserved Structural Elements To Oligomerize and Bind DNA. Mbio, 11, e00448–20.

25. Marko, J.F. and Siggia, E.D. (1994) Fluctuations and Supercoiling of DNA. Science, 265, 506–508.

26. Marko, J.F. (2007) Torque and dynamics of linking number relaxation in stretched supercoiled DNA. Phys. Rev. E, 76, 021926.

27. Danilowicz, C., Hatch, K., Conover, A., Ducas, T., Gunaratne, R., Coljee, V. and Prentiss, M. (2010) Study of force induced melting of dsDNA as a function of length and conformation. J. Phys.: Condens. Matter, 22, 414106.

28. Lipfert, J., Kerssemakers, J.W.J., Jager, T. and Dekker, N.H. (2010) Magnetic torque tweezers: measuring torsional stiffness in DNA and RecA-DNA filaments. Nat. Methods, 7, 977–980.

29. Marko, J.F. and Siggia, E.D. (1995) Stretching DNA. Macromolecules, 28, 8759–8770.

30. Bustamante, C., Marko, J.F., Siggia, E.D. and Smith, S. (1994) Entropic Elasticity of λ-Phage DNA. Science, 265, 1599–1600.

31. Wang, M.D., Yin, H., Landick, R., Gelles, J. and Block, S.M. (1997) Stretching DNA with optical tweezers. Biophys. J., 72, 1335–1346.

32. Bouchiat, C., Wang, M.D., Allemand, J.-F., Strick, T., Block, S.M. and Croquette, V. (1999) Estimating the Persistence Length of a Worm-Like Chain Molecule from Force-Extension Measurements. Biophys. J., 76, 409–413.

33. Nelson, P. (1998) New Measurements of DNA Twist Elasticity. Biophys. J., 74, 2501–2503.

34. Moroz, J.D. and Nelson, P. (1997) Torsional directed walks, entropic elasticity, and DNA twist stiffness. Proc. Natl. Acad. Sci., 94, 14418–14422.

35. Marko, J.F. and Siggia, E.D. (1995) Statistical mechanics of supercoiled DNA. Phys. Rev. E, 52, 2912–2938.

36. Halford, S.E. and Marko, J.F. (2004) How do site-specific DNA-binding proteins find their targets? Nucleic Acids Res., 32, 3040–3052.

37. Graham, J.S., Johnson, R.C. and Marko, J.F. (2011) Concentration-dependent exchange accelerates turnover of proteins bound to double-stranded DNA. Nucleic Acids Res, 39, 2249–2259.

38. Hemphill, W.O., Voong, C.K., Fenske, R., Goodrich, J.A. and Cech, T.R. (2023) Multiple RNA- and DNA-binding proteins exhibit direct transfer of polynucleotides with implications for target-site search. Proc. Natl. Acad. Sci., 120, e2220537120.

39. Kamar, R.I., Banigan, E.J., Erbas, A., Giuntoli, R.D., Cruz, M.O. de la, Johnson, R.C. and Marko, J.F. (2017) Facilitated dissociation of transcription factors from single DNA binding sites. Proc. Natl. Acad. Sci., 114, E3251–E3257.

40. Oehler, S., Amouyal, M., Kolkhof, P., Wilcken-Bergmann, B. von and Müller-Hill, B. (1994) Quality and position of the three lac operators of E. coli define efficiency of repression. EMBO J., 13, 3348–3355.

41. Hartmann, G., Honikel, K.O., Knüsel, F. and Nüesch, J. (1967) The specific inhibition of the DNA-directed RNA synthesis by rifamycin. Biochim. Biophys. Acta (BBA) - Nucleic Acids Protein Synth., 145, 843–844.

42. Zhou, B., Schrader, J.M., Kalogeraki, V.S., Abeliuk, E., Dinh, C.B., Pham, J.Q., Cui, Z.Z., Dill, D.L., McAdams, H.H. and Shapiro, L. (2015) The Global Regulatory Architecture of Transcription during the Caulobacter Cell Cycle. Plos Genet, 11, e1004831.

43. Lalanne, J.-B., Taggart, J.C., Guo, M.S., Herzel, L., Schieler, A. and Li, G.-W. (2018) Evolutionary Convergence of Pathway-Specific Enzyme Expression Stoichiometry. Cell, 173, 749–761.e38.

44. Li, G.-W., Burkhardt, D., Gross, C. and Weissman, J.S. (2014) Quantifying Absolute Protein Synthesis Rates Reveals Principles Underlying Allocation of Cellular Resources. Cell, 157, 624–635.

45. Yan, J., Kawamura, R. and Marko, J.F. (2005) Statistics of loop formation along double helix DNAs. Phys. Rev. E, 71, 061905.

46. Dehal, P.S., Joachimiak, M.P., Price, M.N., Bates, J.T., Baumohl, J.K., Chivian, D., Friedland, G.D., Huang, K.H., Keller, K., Novichkov, P.S., et al. (2010) MicrobesOnline: an integrated portal for comparative and functional genomics. Nucleic Acids Res, 38, D396– D400.

